# Kel1 is a phosphorylation-regulated noise suppressor of the pheromone signaling pathway

**DOI:** 10.1101/2021.05.19.443414

**Authors:** Ignacio Garcia, Sara Orellana-Muñoz, Lucía Ramos-Alonso, Aram N. Andersen, Christine Zimmermann, Jens Eriksson, Stig Ove Bøe, Petra Kaferle, Manolis Papamichos-Chronakis, Pierre Chymkowitch, Jorrit M. Enserink

**Affiliations:** Department of Molecular Cell Biology, Institute for Cancer Research, The Norwegian Radium Hospital, Montebello, 0379 Oslo, Norway; Centre for Cancer Cell Reprogramming, Institute of Clinical Medicine, Faculty of Medicine, University of Oslo, 0318 Oslo, Norway; Section for Biochemistry and Molecular Biology, Faculty of Mathematics and Natural Sciences, University of Oslo, 0037 Oslo, Norway; Department of Microbiology, Oslo University Hospital, 0372 Oslo, Norway; Institute for Virology, University Medical Center of the Johannes Gutenberg-University, 55131 Mainz, Germany; Department of Medical Biochemistry and Microbiology, Uppsala University, 752 37 Uppsala, Sweden; Institut Curie, PSL Research University, CNRS, UMR3664, Sorbonne Universities, France; Department of Molecular Physiology and Cell Signalling Institute of Systems, Molecular and Integrative Biology University of Liverpool, L69 7BE Liverpool, UK

## Abstract

Mechanisms have evolved that allow cells to detect signals and generate an appropriate response. The accuracy of these responses relies on the ability of cells to discriminate between signal and noise. How cells filter noise in signaling pathways is not well understood. Here, we analyze noise suppression in the yeast pheromone signaling pathway and show that the poorly characterized protein Kel1 serves as a major noise suppressor and prevents cell death. At the molecular level, Kel1 prevents spontaneous activation of the pheromone response by inhibiting membrane recruitment of Ste5 and Far1. Only a hypophosphorylated form of Kel1 suppresses signaling, reduces noise and prevents pheromone-associated cell death, and our data indicate that the MAPK Fus3 contributes to Kel1 phosphorylation. Taken together, Kel1 serves as a phospho-regulated suppressor of the pheromone pathway to reduce noise, inhibit spontaneous activation of the pathway, regulate mating efficiency, and to prevent pheromone-associated cell death.

## Introduction

A crucial aspect of any organism’s well-being is the ability of cells to respond to changes in their internal and external milieu. Accurate signal-noise discrimination is particularly important during conditions that threaten cellular homeostasis, or when a given signal triggers cellular commitment, such as differentiation. Low levels of noise within a population of cells may be beneficial under certain conditions, by allowing a fraction of cells to survive a dramatic change in environmental conditions (Kaern et al., 2005). However, high noise levels may be detrimental to cellular fitness and has been evolutionarily minimized (Balazsi et al., 2011; Lehner, 2008; Metzger et al., 2015; Wang and Zhang, 2011). Noise has been best studied at the level of gene expression, where it is often referred to as the stochastic variation in the protein expression level of a gene among isogenic cells in a homogenous environment (Raser and O’Shea, 2005; Wang and Zhang, 2011). Gene expression noise can arise from intrinsic and extrinsic variations (Raser and O’Shea, 2005). Intrinsic noise is caused by inherent stochastic events in biochemical processes that can occur at various levels during gene expression, such as transcriptional initiation, mRNA degradation, translational initiation, and protein degradation, as well as during signal transduction (Raser and O’Shea, 2005). Extrinsic noise is caused by differences among cells, either in their local environment or the concentration or activity of any factor that influences gene expression (Raser and O’Shea, 2005; Volfson et al., 2006), such as age, cell cycle stage, metabolic state, and the number and quality of proteins and organelles distributed to the mother and daughter cell during cell division.

The yeast mating pathway is a model for signal transduction and decision-making (Alvaro and Thorner, 2016; Paliwal et al., 2007). In haploid yeast cells, this pathway detects and transmits a pheromone signal emitted by cells of the opposite mating type to induce a mating response (Fig. 1A). Pathway activation results in cell cycle arrest, activation of a transcriptional program and cell wall remodeling to execute the morphological changes required to mate (Alvaro and Thorner, 2016). Given the potential fitness cost associated with inappropriate activation of this pathway (Banderas et al., 2016), signaling occurs in a switch-like manner with high precision and overall low intrinsic noise (Dixit et al., 2014; Malleshaiah et al., 2010). The pathway is activated by binding of pheromone to its G protein-coupled target receptor, resulting in dissociation of the heterotrimeric G protein into the α-subunit (Gpa1) and a Gβγ heterodimer (Ste4–Ste18)(Fig. 1A). The Gβγ complex recruits Far1 and the scaffold protein Ste5 to the plasma membrane. Membrane localization of Ste5 results in recruitment and activation of a MAPK module composed of Ste11, Ste7 and the MAPKs Fus3 and Kss1 (Alvaro and Thorner, 2016). Fus3 then phosphorylates and stimulates the transcription factor Ste12 (Elion et al., 1993), which induces a transcriptional program required for efficient mating. Membrane-localized Far1 mediates polarized growth by recruiting the guanine nucleotide exchange factor Cdc24, which stimulates Cdc42 to induce formation of a mating projection (shmoo) (Butty et al., 1998).

**Figure 1.**
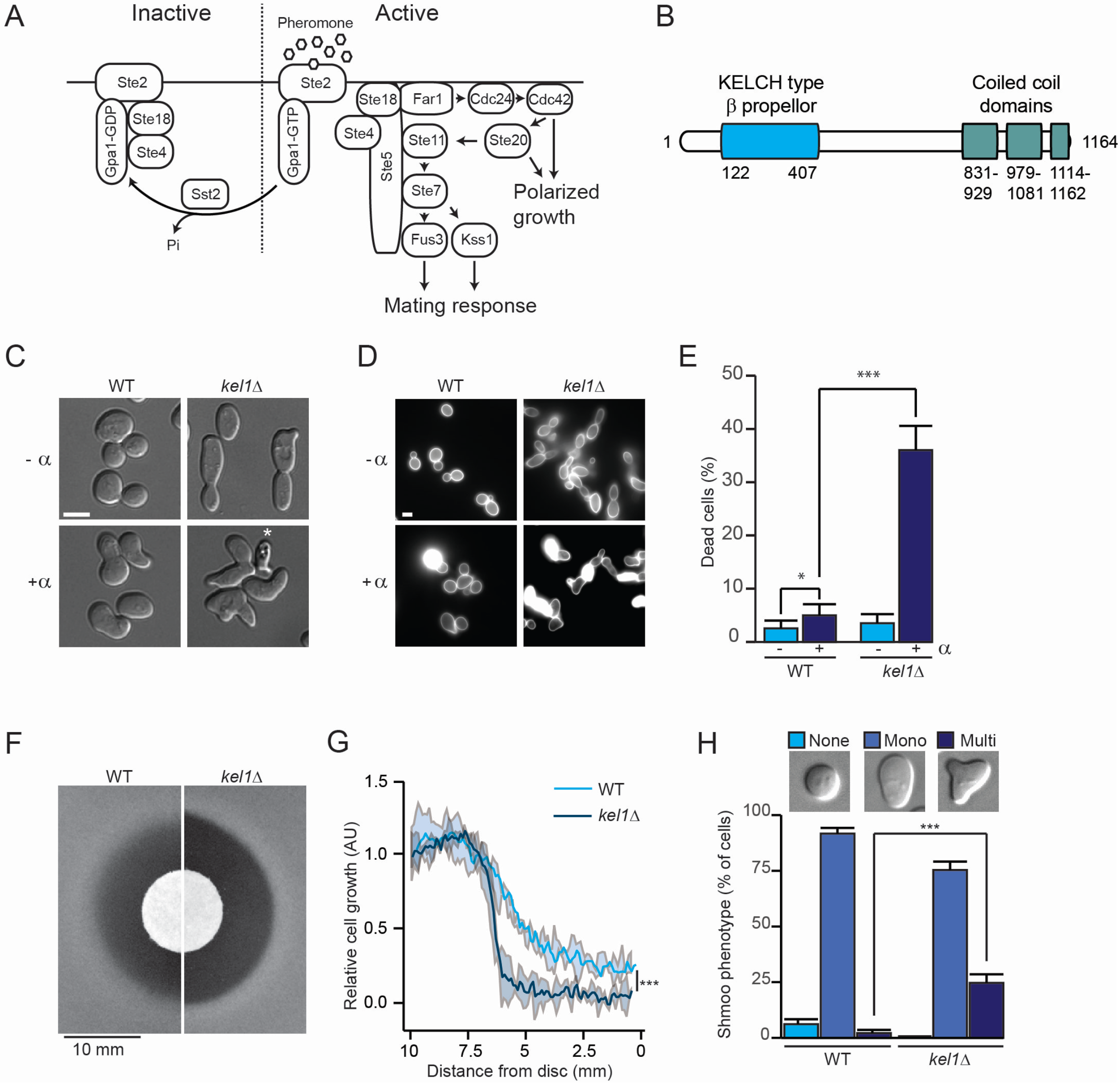
Kel1 prevents formation of multiple shmoos and suppresses cell death in the presence of pheromone. ***A,*** Overview of the pheromone response pathway. ***B***, Protein domains in Kel1. ***C***, Morphological defects of *kel1*Δ mutants. DIC images of WT and *kel1*Δ cells in absence or presence of pheromone. Asterisk, dead cell. Bar, 5 µM. ***D***, Kel1 prevents cell death. Fluorescence microscope images of WT and *kel1*Δ cells incubated in absence or presence of pheromone. Cells were stained with FITC (0.01 mg/ml) for 5 min to visualize cell death. Bar, 5 µM. ***E***, Percentages of dead cells. Bars: Mean of 5 independent experiments (average of 625 cells per strain and condition); Error bars: Bootstrapped-standard deviations. Statistical significance was calculated using two-proportions z-tests, * 0.01<p<0.05; *** p<0.001. ***F***, Pheromone sensitivity assay. Cells were plated in a top layer of agar on which a sterile filter was placed containing 15 μg of pheromone. ***G***, Densitometric quantification of relative cell growth across the plates shown in (F)(n=3). Statistical significance of the density at the region closer to the disc was calculated using t-test, standard deviation p<0.001. ***H***, Quantification of the number of shmoos per cell in cultures of WT and *kel1*Δ strains treated with pheromone (132 cells per strain and condition from 5 independent samples). Bars, mean from three independent experiments; Error bars, bootstrapped-standard deviations. Statistical significance was calculated using two-proportions z-tests, *** p<0.001.

Feedback loops have been identified that improve transmission of information and that help switch off the pathway. An example of positive feedback is the increased expression of *FUS3* that occurs upon activation of Ste12 by Fus3 (Roberts et al., 2000). Fus3 also provides negative feedback to dampen the response and help switch off the mating response through inhibition of Ste5 recruitment (Choudhury et al., 2018; Repetto et al., 2018; Yu et al., 2008). This is physiologically relevant, because hyperactivation of the mating pathway, or an attempt to mate in absence of a partner, can lead to cell death (Severin and Hyman, 2002; Zhang et al., 2006). Another negative regulator is Sst2, which is a Regulator of G protein Signaling (RGS) that inhibits signaling by Gpa1 (Apanovitch et al., 1998) and which mediates desensitization of cells after pheromone treatment (Dohlman et al., 1995). Sst2 also functions as a noise suppressor (Dixit et al., 2014). The dominant source of variation in the pheromone pathway is thought to be extrinsic noise (Colman-Lerner et al., 2005), and mutant cells either lacking *SST2* or expressing a mutant form of Gpa1 that is resistant to the GAP activity of Sst2 show increased levels of noise (Dixit et al., 2014). Noise suppression is required for proper gradient detection and morphogenesis, and one potential physiological consequence of elevated noise in mutant cells is reduced mating efficiency (Dixit et al., 2014).

Despite the fact that pheromone signaling has been studied intensively during the past decades, there are still several genes with poorly characterized functions in the pathway. One such gene is *KEL1*, which encodes a 131 kDa protein consisting of a KELCH propeller three coiled-coil domains (Fig. 1B). Kel1 was first identified in a screen for genes whose overexpression relieved the mating defect caused by activated alleles of *PKC1* (Philips and Herskowitz, 1998). *kel1Δ* mutants are elongated and heterogeneous in shape and have a defect in cell fusion. Kel1 localizes to sites of polarized growth and forms a ternary complex with the cell fusion regulator Fus2 and activated Cdc42 during mating (Smith and Rose, 2016). Kel1 also interacts with formins to regulate the assembly of actin cables (Gould et al., 2014). These findings suggest that Kel1 may serve as a hub during the pheromone response, but how and to which extent Kel1 controls the pheromone response remains unknown.

Here, we characterized the function of Kel1 in the pheromone response. We present data showing that Kel1 is regulated by phosphorylation, suppresses spontaneous activation of the pheromone pathway, has a major role in filtering noise in the pheromone pathway, and that it prevents pheromone-induced cell death.

## Results

### Kel1 is important for the pheromone response

During the course of our experiments, we serendipitously observed that approximately 35% of *kel1*Δ mutant cells died upon treatment with pheromone (Fig. 1C-E). Careful analysis revealed that also a small number of wild-type (WT) cells died after pheromone treatment (Fig. 1E), consistent with previous findings (Severin and Hyman, 2002). The importance of Kel1 in preventing cell death was reflected in pheromone halo assays, where we observed a 30% cell-density reduction in the *kel1*Δ strain compared with the WT strain inside the halo, where the concentration of pheromone is highest (Fig. 1F,G). Although the area of the halo was not significantly different, the interface was sharper (Fig. 1F,G), consistent with a role for Kel1 in the adaptive response to pheromone. Indeed, while *kel1*Δ mutants had significant morphological defects during vegetative growth, such as elongated buds and misshapen cells (Suppl. Fig. S1A,B), they were also 15 times more prone than WT cells to initiate more than one shmoo during pheromone treatment (Fig. 1H).

Together, these data indicate that Kel1 prevents pheromone-associated cell death.

### Kel1 activity is regulated by phosphorylation

Phosphoproteome studies have detected at least 50 phosphorylated amino acids in Kel1, a substantial number of which are [S/T-P] sites (https://thebiogrid.org/36592/protein; Fig. 2A), which are typically targeted by proline-directed kinases such as CDKs and MAPKs. Interestingly, λ-phosphatase treatment of Kel1 resulted in increased mobility on Phos-tag SDS-PAGE, confirming that Kel1 exists as a phosphoprotein *in vivo* (Fig. 2B, lane 2 and 3; quantified in Suppl. Fig. S1C). Kel1 phosphorylation further increased upon pheromone treatment (Fig. 2B, lane 4), consistent with a previous phosphoproteomics report (Li et al., 2007). We hypothesized that proline-directed phosphorylation might regulate Kel1 and constructed two mutant alleles in which all [S/T-P] sites were substituted with either alanine or aspartate residues (*kel1-ala* and *kel1-asp*, respectively, see STAR Methods). These alleles were fused to a C-terminal FLAG tag and integrated into the *KEL1* locus under control of the endogenous promoter. Phos-tag gel electrophoresis of immunoprecipitated proteins revealed that Kel1-ala migrated faster than wild-type Kel1, while phosphatase treatment did not substantially increase its mobility (Fig. 2B, lane 6 and 7). Furthermore, treatment with pheromone did not increase phosphorylation of this mutant (Fig. 2B, lane 8 and 9). We conclude that Kel1 is phosphorylated on one or several [S/T-P] sites *in vivo*.

**Figure 2.**
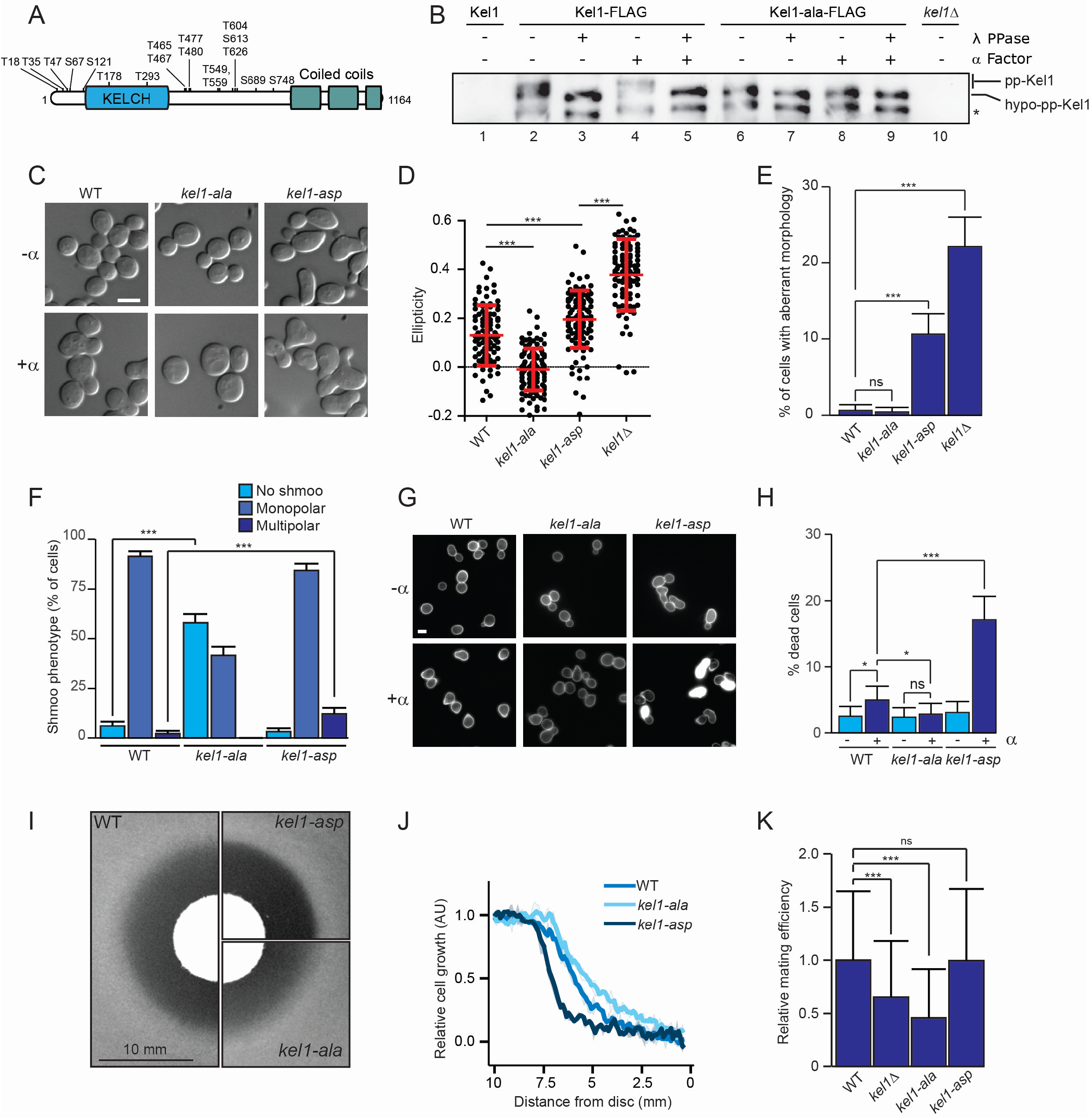
Kel1 is regulated by phosphorylation. ***A***, Potential S/T-P phosphorylation sites. ***B***, Kel1 exists as a phosphoprotein *in vivo*. Cells were either left untreated or were treated with pheromone for 2h, after which Kel1-FLAG and Kel1-ala-FLAG were purified and treated with λ phosphatase, followed by Phos-tag gel electrophoresis and western blotting using FLAG antibodies. Untagged *KEL1* and *kel1Δ* mutant strains were used as negative controls. Asterisk indicates a proteolytically processed/degraded form of Kel1. ***C***, Cell morphology is controlled by Kel1 phosphorylation. DIC images of representative WT, *kel1-ala* and *kel1-asp* cells in absence or presence of pheromone. Bar, 5 µM. ***D,*** Quantification of bud ellipticity of the experiment shown in (C). Ellipticity was determined as in Fig. S1A. Mean and standard deviations for each strain are shown as red bars. Statistical significance was calculated using t-test, *** p<0.001; n=3, analyzing at least 300 cells each (note that only 100 cells were randomly plotted for clarity). ***E,*** Quantification of cells with aberrant morphology in absence of pheromone (Figs. 1C and 2C). Bars, mean from three independent experiments (at least of at least 217 cells each); error bars, bootstrapped-standard deviation. Statistical significance was calculated using two-proportions z-tests, *** p<0.001. ***F,*** Efficient shmoo formation requires Kel1 phosphorylation. Quantification of the number of shmoos per cell (132 cells per strain and per condition from 5 independent experiments) was performed as described in Fig. 1H, *** p<0.001. ***G,*** Increased death of *kel1-asp* mutants after treatment with pheromone, visualized as in Fig. 1D. Bar, 5 µM. ***H,*** Quantification of cell death, as described in Fig. 1E, * 0.01<p<0.05. ***I,*** Pheromone filter assay, performed as described in Fig. 1F. ***J,*** Densitometric quantification of the experiment shown in (I), including two additional independent experiments (n=3). ***K,*** *kel1* mutants have mating defects. Mating efficiency assay was performed as described in STAR Methods. Bars, mean from three independent experiments; Error bars, bootstrapped-standard deviation, *** p<0.001.

We noticed that during vegetative growth the buds of WT cells were slightly oblong, whereas *kel1-ala* buds were almost perfectly spherical (Fig. 2C,D). *kel1-asp* buds were significantly more elongated than WT buds, although not as much as those of *kel1*Δ cells (Fig. 2C,D). Both *kel1-asp* and *kel1Δ* mutant cells frequently showed aberrant morphology (Fig. 2E). More importantly, upon pheromone treatment more than half of the *kel1-ala* mutant cells failed to form a shmoo (Fig. 2C,F), and those cells that did form a shmoo always generated a single, highly spherical shmoo (see below). In contrast, *kel1-asp* mutant cells readily formed shmoos and were more likely to form multiple shmoos than WT cells (Fig. 2F). Thus, the phenotype of *kel1-asp* cells resembles that of *kel1*Δ mutant cells, albeit more modest. Pheromone-associated cell death was also higher in *kel1-asp* mutants than in WT cells, whereas in *kel1-ala* mutants it was lower (Fig. 2G,H). This was reflected in the pheromone halo assay, which showed that the cell-growth interphase was sharper in the *kel1-asp* strain and more diffuse in the *kel1-ala* mutant compared to WT (Fig. 2I,J). Finally, the mating capacity of *kel1*Δ and *kel1-ala* mutants was significantly reduced (Fig. 2K).

These results indicate that phosphorylation of Kel1 is important for regulation of the mating process. Given that *kel1-ala* mutants have a phenotype opposite to that of phospho-mimicking *kel1-asp* mutants, and that the phenotype of *kel1-asp* cells generally resembles that of *kel1*Δ cells (although milder), we conclude that (i) hypophosphorylated Kel1 suppresses the pheromone response, and (ii) phosphorylation of certain [S/T-P] sites relieves the inhibitory effect of Kel1 on the pheromone response.

### Kel1 acts downstream of Sst2 during the mating response

To identify pathways associated with phosphorylation of Kel1 we performed a genome-wide synthetic genetic array (SGA) screen (Baryshnikova et al., 2010; Tong et al., 2001). Since *kel1-asp* mutants have a loss-of-function phenotype that generally resembles that of the *kel1*Δ strain, we decided to use only the *kel1*Δ and *kel1-ala* strains as query mutants. These mutants were crossed into the library of non-essential knock-out genes and the Decreased Abundance by mRNA Perturbation (DAmP) library (Breslow et al., 2008; Giaever et al., 2002). For all genetic interactions we calculated genetic interaction scores (STAR Methods). We focused on a subset of genes that showed differential genetic interactions with the *kel1*Δ and *kel1-ala* mutations, using a strict cut-off with differences of at least three standard deviations from the mean (Fig. 3A, Suppl. Fig. S1D, and Suppl. Dataset S1). To find patterns in this gene list, we studied the reported phenotypes for these genes using information gathered from the *Saccharomyces* Genome Database. The most highly overrepresented phenotype was *“Resistance to enzymatic treatment”* (Fig. 3B). Mutations in genes associated with this phenotype often lead to cell wall defects, resulting in increased sensitivity to cell wall-lytic enzymes. This is consistent with our finding that *kel1*Δ mutants frequently undergo cell death during pheromone-induced cell wall remodeling. The second most overrepresented phenotype was *“Mating response”*. This is likely an underestimate, because many pheromone-associated genes are absent from SGA screens due to severe mating defects.

**Figure 3.**
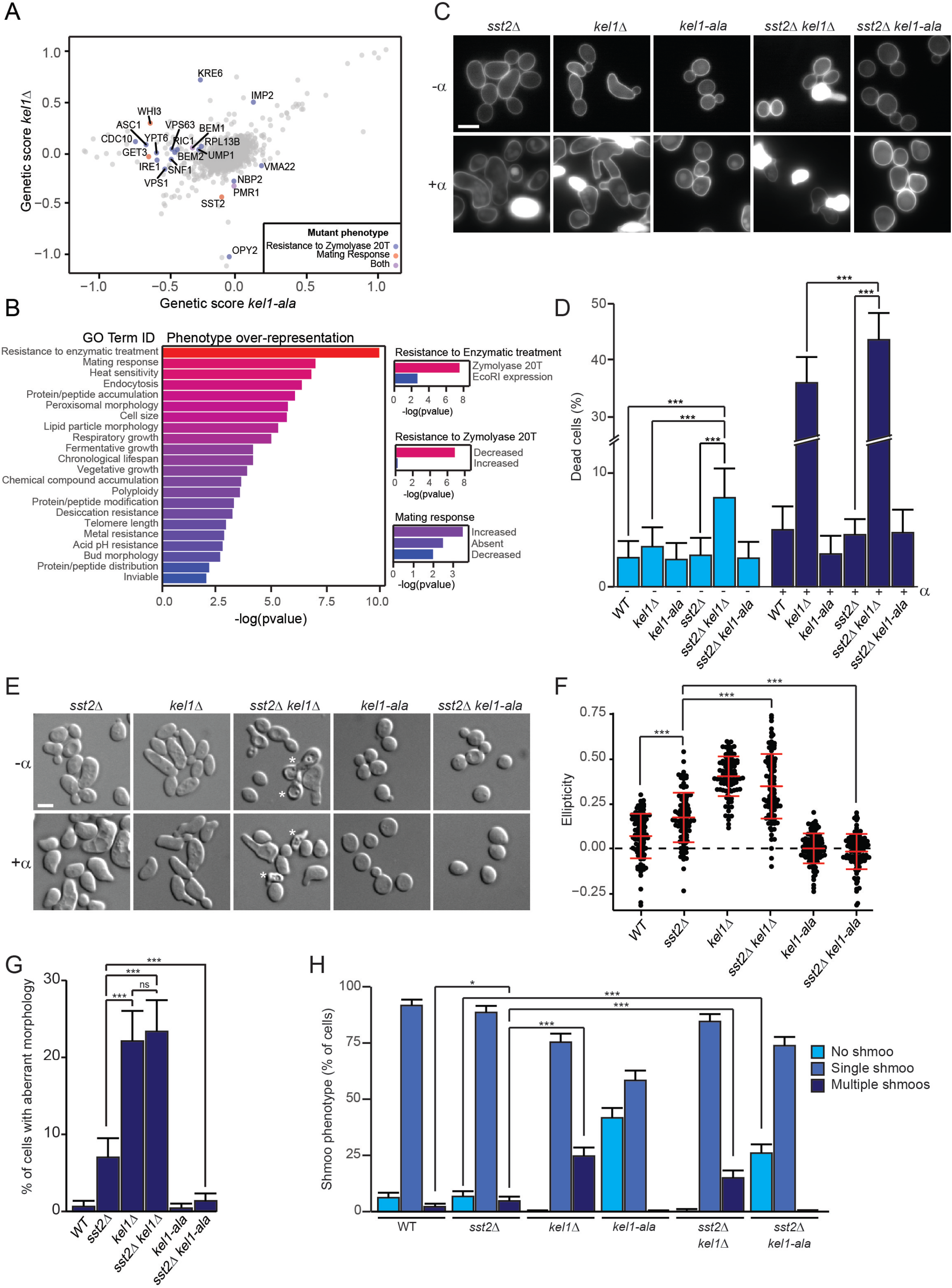
*KEL1* acts downstream of Sst2 in the pheromone response. ***A,*** Results of the SGA screen. Dots represent genes, coordinates represent genetic scores observed with the *kel1*Δ versus the *kel1-ala* strain. Colored dots indicate genes with reported phenotypes in mating and/or resistance to Zymolyase that show differential genetic scores in the *kel1* mutants. ***B,*** Gene ontology enrichment analysis of the genes showing differential genetic scores with *kel1*Δ and *kel1-ala*. Calculation of enrichment is described in STAR Methods. ***C,*** Loss of *SST2* enhances the toxicity of pheromone in *kel1Δ* mutants. The experiment was performed as described in Figure 1D. Bar, 5 µM. ***D,*** Quantification of cell death, described as in Figure 1E, *** p<0.001 (132 cells per strain and per condition from 5 independent experiments). ***E,*** *kel1-ala* suppresses the morphological defects of the *sst2*Δ mutant. Microscopic images of the indicated strains in absence or presence of pheromone. Asterisk: Dead cells. Bar, 5 µM. ***F,*** Quantification of bud ellipticity of untreated cells in the experiment shown in (E), including two additional independent experiments (300 cells each, only 100 cells were randomly plotted for clarity). The analysis and statistics were performed as described in Figure 2D. ***G,*** Quantification of cells with aberrant morphology in untreated cells (E) was performed as in Fig. 2E (at least of at least 217 cells each), *** p<0.001. ***H,*** Expression of *kel1-ala* inhibits shmoo formation of *sst2*Δ mutant cells. Quantification of the number of shmoos per cell (132 cells per strain and per condition from 5 independent experiments) was performed as in Fig. 1H, * 0.01<p<0.05; *** p<0.001.

We were intrigued by one mating response gene in particular, i.e. *SST2*. Loss of *SST2* substantially reduced the fitness of *kel1*Δ mutants, but did not appear to have a strong effect on the fitness of *kel1-ala* mutants (Fig. 3A). Even in absence of pheromone approximately 8% of the cells in *sst2*Δ *kel1*Δ cultures were dead (Fig. 3C,D). Pheromone treatment strongly increased cell death of *kel1*Δ mutants, and although cell death in *sst2*Δ cultures was similar to WT (Fig. 3D), it was significantly higher in *sst2*Δ *kel1*Δ double mutants than in either single mutant (Fig. 3C,D).

We next studied the cellular morphology of the mutants in presence and absence of pheromone. 7% of vegetatively growing *sst2*Δ cells showed aberrant morphology, and the buds of *sst2*Δ mutants were generally more elongated than those of WT cells, but to a lesser extent than *kel1*Δ mutants (Fig. 3E-G). Pheromone treatment resulted in a small but significant increase in the number of *sst2*Δ cells with multiple projections (Fig. 3H). The morphology of the *sst2*Δ *kel1*Δ mutant resembled that of *kel1*Δ (Fig. 3F,G), and pheromone treatment of *sst2*Δ *kel1*Δ mutants did not further increase the number of cells with multiple projections compared with the *kel1*Δ single mutant (Fig. 3E,H). However, the *kel1-ala* mutation completely suppressed the morphological defects and inhibited shmoo formation of *sst2*Δ mutants (Fig. 3E-H).

Taken together, these data show that Kel1 and Sst2 regulate cell morphogenesis both during vegetative growth and in the presence of pheromone, with Kel1 having a dominant role, and that Kel1 functions downstream of Sst2 in this process.

### Kel1 physically interacts with pheromone pathway components and may be phosphorylated by Fus3

To better understand how Kel1 is regulated, we immunopurified Kel1 from untreated and pheromone-treated cells and identified interaction partners using mass-spectrometry (Supplemental Dataset S2). We identified several known Kel1 interactors, thus validating the approach (Suppl. Fig. S1E). We looked for proteins that differentially interacted with Kel1 depending on pheromone treatment, and observed a significant enrichment of proteins with functions in the mating response (Fig. 4A), including Far1, Sst2, Fig1, the pheromone receptor Ste2 and the MAPK Fus3. We validated the pheromone-induced interaction between Kel1 and Ste2 and Fus3 in co-immunoprecipitation experiments (Fig. 4B). The interaction with the proline-directed kinase Fus3 piqued our interest, since Kel1 is phosphorylated on [S/T-P] sites in vivo. We evaluated Kel1 phosphorylation using Phos-tag gels in WT cells, in a *fus3*Δ *kss1*Δ double mutant, and in mutants lacking the CDK Pho85, which we included for comparison. In untreated cells, there was no difference in mobility of Kel1 between WT cells and *fus3*Δ *kss1*Δ double mutants (Fig. 4C, lane 1 and 2). Somewhat unexpectedly, Kel1 phosphorylation appeared to be reduced in vegetatively grown *pho85Δ* mutants (lane 3), which will be studied in more detail in future studies. More importantly, in pheromone-treated cells, phosphorylation of Kel1 was partially reduced in the *fus3*Δ *kss1*Δ double mutant (Fig. 4C, lane 7, compare with lane 6; Suppl. Fig. S1G). It should be noted that, compared to Kel1-ala, a substantial fraction of Kel1 remained phosphorylated in *fus3*Δ *kss1*Δ double mutant cells (compare lane 7 to lanes 9 and 10), indicating that Kel1 is also phosphorylated by other kinases (see Discussion). Because the possible number of combinations of kinase mutations is relatively large, this will be the focus of future work.

**Figure 4.**
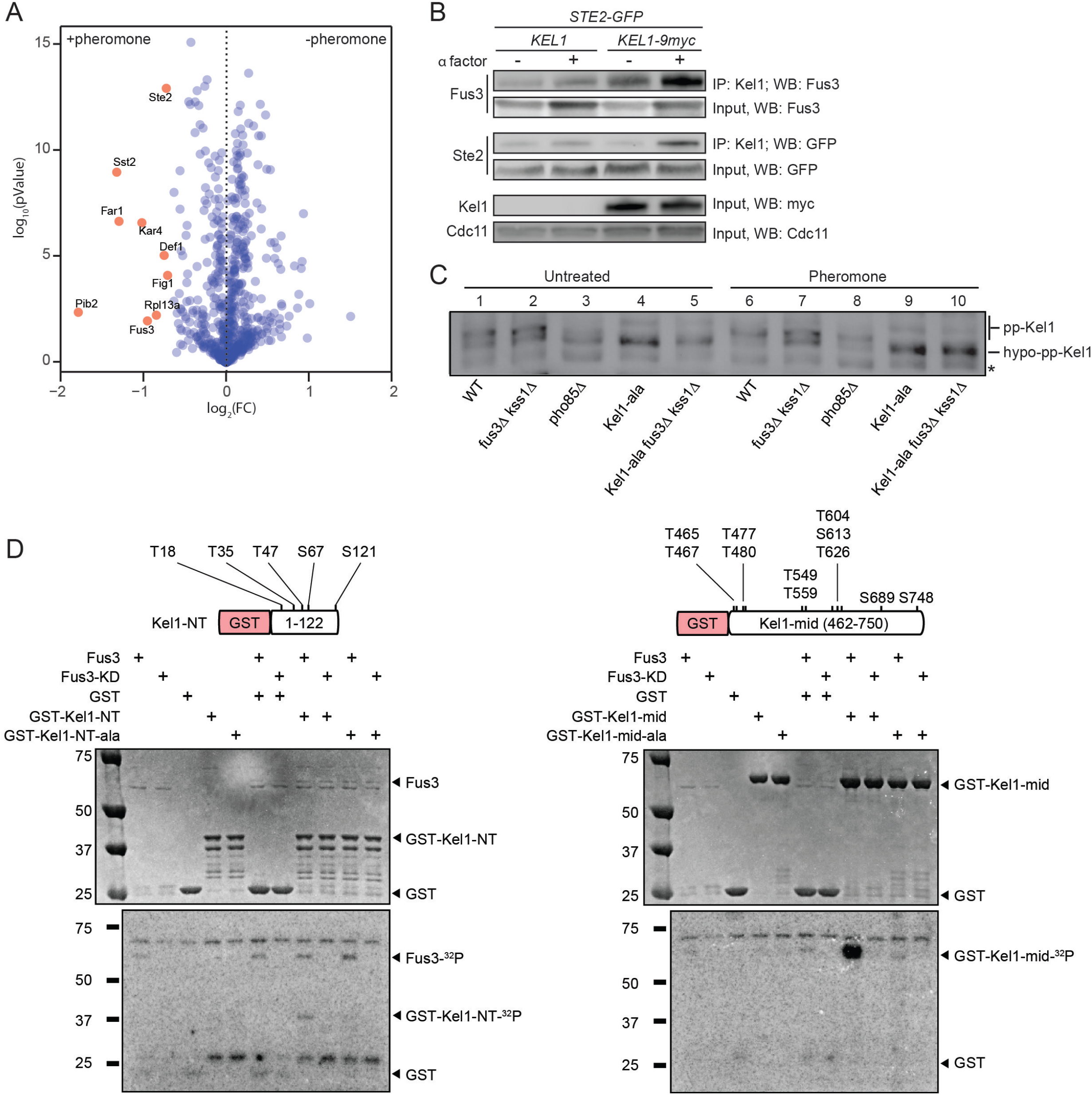
Kel1 binds multiple components of the mating response pathway in response to pheromone. ***A,*** Volcano plot of IP-MS experiments using Myc-tagged Kel1 in absence or presence of pheromone. Enrichment of interaction partners was determined by quantitative MS. Binding partners enriched >1.6-fold after treatment with pheromone (p-value<0.013) are shown in red. ***B,*** Pheromone-induced physical interactions between Kel1-Ste2 and Kel1-Fus3. Myc-tagged Kel1 was immunoprecipitated from cells expressing Ste2-GFP, after which cell lysates were analyzed by western blotting using GFP- and Fus3-specific antibodies. ***C,*** Efficient phosphorylation of Kel1 during pheromone treatment requires Fus3/Kss1. Experiment was described as in Figure 2B. ***C,*** Fus3 can phosphorylate Kel1 *in vitro*. Two recombinant Kel1 fragments covering all but two potential S/T-P sites, their alanine substitutions, as well as plain GST were subjected to *in vitro* kinase assays with recombinant Fus3 or kinase-dead Fus3 (Fus3-KD) as described in STAR Methods.

We then asked whether Fus3 can directly phosphorylate Kel1 in vitro. Recombinant full-length Kel1 was insoluble and therefore we purified two Kel1 fragments, Kel1-NT (1-122) and Kel1-mid (462-750), which together cover all but two S/T-P sites (T178 and T293). Interestingly, Fus3, but not kinase-dead Fus3, efficiently phosphorylated Kel1-mid, and to a much lesser degree also Kel1-NT, and it failed to phosphorylate fragments with alanine substitutions (Fig. 4D).

Taken together, these data suggest that Fus3 is at least partially responsible for Kel1 phosphorylation, although other kinases likely also contribute to this process.

### Kel1 prevents spontaneous recruitment of Ste5 and Far1 to inhibit formation of mating projections in absence of pheromone

Vegetatively growing *kel1Δ* mutant cells often formed elongated structures that resemble shmoos, and therefore we monitored membrane localization of Ste5 and Far1, which are markers for formation of mating projections (Nern and Arkowitz, 1999). Strikingly, even in absence of pheromone, we observed that NeonGreen (mNG)-tagged Ste5 and Far1 were spontaneously recruited to patches at the cell cortex both in *kel1Δ* mutants and in *kel1-asp* mutant cells (Fig. 5A,B). This was not observed in WT cells or in *kel1-ala* mutants. Whereas pheromone treatment resulted in localization of Ste5-mNG and Far-mNG at the cortex of WT cells, this was undetectable in the *kel1-ala* mutant (Fig. 5A-C). These findings show that the polarized structures in vegetatively growing *kel1*Δ and *kel1-asp* mutants are indeed spontaneous shmoos, and suggest that Kel1 suppresses spontaneous signaling through the pheromone pathway. Therefore, we analyzed the mRNA levels of two genes known to be upregulated by the pheromone pathway, i.e. *FUS3* and *STE2* (Oehlen et al., 1996). Although there was no significant difference in *FUS3* and *STE2* mRNA levels between populations of WT cells and any of the *kel1* mutants after pheromone treatment, *kel1*Δ mutant populations showed a significant increase in mRNA levels in absence of pheromone (Fig. 5D), confirming that Kel1 suppresses spontaneous pathway activity.

**Figure 5.**
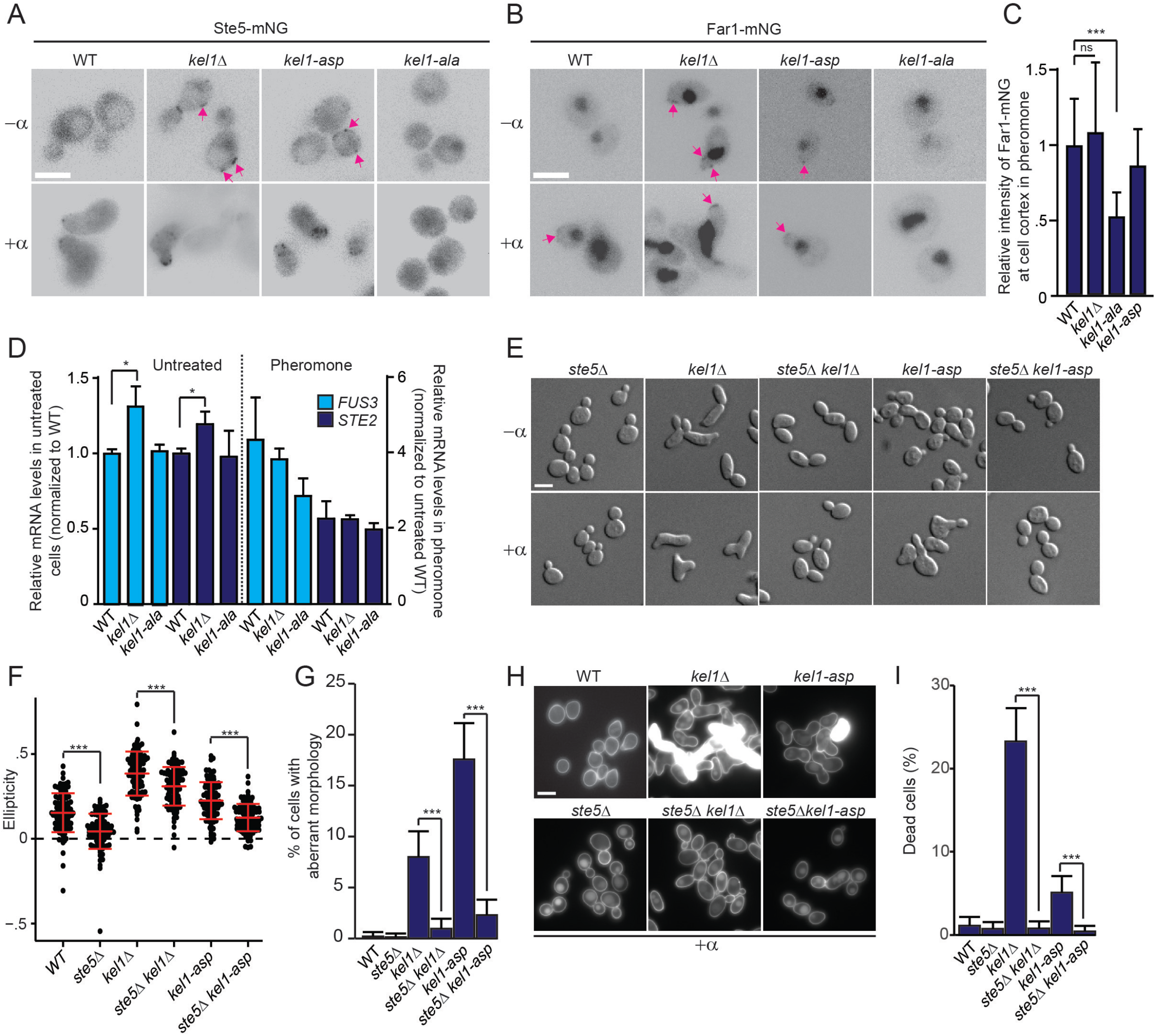
Kel1 functions upstream of Ste5 to prevent spontaneous formation of shmoos. ***A,*** Kel1 prevents ectopic recruitment of Ste5 in absence of pheromone. mNeonGreen-tagged Ste5 (Ste5-mNG) was imaged in living cells in absence or presence of pheromone. Arrows: Cortex-localized Ste5. Bar, 5 µM. ***B,*** Cortical recruitment of Far1-mNG was studied as in (A). Bar, 5 µM. ***C,*** Quantification of the overall intensity of Far1-mNG (B) in presence of pheromone. Bars represent the mean of the relative intensity and the error bars the standard deviation. Statistical significance was calculated using t-test, *** p<0.001. ***D,*** Kel1 prevents spontaneous expression of pheromone-induced mRNAs. *FUS3* and *STE2* mRNA levels were analyzed in WT and *kel1* mutants in absence and presence of pheromone. mRNA levels were normalized to WT. Error bars: Standard deviation. Statistical significance was calculated using t-test, * 0.01<p<0.05; three independent experiments. ***E,*** Deletion of *STE5* rescues *kel1*Δ phenotypes. The indicated strains were incubated in absence or presence of pheromone before imaging. Bar, 5 µM. ***F,*** Quantification of bud ellipticity of untreated cells in the experiment shown in (E). Mean and standard deviations for each strain are shown as red bars. Statistical significance was calculated using t-test, *** p<0.001 ***G,*** Quantification of cells with aberrant morphology in absence of pheromone of the experiment shown in (E). Analysis as in Figure 2E, *** p<0.001. ***H,*** Deletion of *STE5* prevents cell death in the *kel1*Δ and *kel1-asp* mutants. Cell death was visualized as in Figure 1D. WT image was taken from an independent experiment. Bar, 5 µM. ***I,*** Quantification of cell death of the experiment shown in (H). Analysis as in Figure 1E, ** 0.001<p<0.01.

If Kel1 indeed inhibits spontaneous signaling through the pheromone pathway by preventing the accumulation of Ste5 at the cell cortex, then deletion of *STE5* should reverse the phenotype of *kel1*Δ and *kel-asp* mutants. The buds of vegetatively growing *ste5*Δ mutants were significantly more spherical than those of WT cells (Fig. 5E,F), and deletion of *STE5* also significantly reduced the aberrant bud and cell morphology phenotypes of *kel1*Δ and *kel1-asp* mutant cells (Fig. 5E-G). Importantly, deletion of *STE5* almost completely rescued pheromone treatment-induced lethality in *kel1*Δ and *kel1-asp* mutants (Fig. 5H,I). It should be noted that the absence of Ste5 has multiple effects on the cell, i.e. it not only diminishes expression of multiple pathway components, but it also prevents the recruitment and activation of pathways components that are expressed. Nonetheless, these data show that Kel1 suppresses spontaneous activation of the pheromone pathway by preventing aberrant recruitment of Ste5 to the cell cortex.

### Hypophosphorylated Kel1 cooperates with Sst2 to prevent spontaneous pheromone signaling and to dampen the pheromone response

Spontaneous activation of the pheromone pathway has been observed previously and was found to originate downstream of the receptor at the level of the G protein (Siekhaus and Drubin, 2003), which we confirmed (Suppl. Fig. S1F). Mechanisms exist that suppress spontaneous signaling; for instance, RGS proteins like Sst2 inhibit unscheduled G protein signaling (Siekhaus and Drubin, 2003). Given our findings that Kel1 acts downstream of Sst2 but at/upstream of Ste5, and that the *kel1-ala* mutant suppresses the morphological defects observed in *sst2*Δ mutant cells, we hypothesized that Kel1 might cooperate with Sst2 to inhibit spontaneous signaling. We made use of a dual reporter system based on expression of GFP under control of the pheromone pathway-sensitive *FUS1* promoter and mCherry under control of the pheromone pathway-independent *ADH1* promoter; the ratio of GFP/mCherry is a measure of signaling pathway activity at single-cell resolution (Dixit et al., 2014). Interestingly, single-cell microscopy revealed that in absence of pheromone the GFP/mCherry ratio was significantly higher in *kel1*Δ mutants than in WT cells (“0 min” in Fig. 6A; timecourse in Fig. 6B; quantification in Fig. 6C), confirming our finding that Kel1 serves as a major signal suppressor in absence of stimulus. We then analyzed the relative contributions of Sst2 and Kel1 in suppressing spontaneous activity of the signaling pathway. In absence of pheromone the basal level of the signal was higher in the *sst2*Δ mutant than in the WT strain, and was similar to that of *kel1*Δ mutants (Fig. 6A-C). However, *sst2*Δ *kel1*Δ double mutants had significantly higher basal signaling levels than either single mutant, whereas the *kel1-ala* mutation suppressed spontaneous pathway activity in the *sst2*Δ mutant to levels that were not significantly different from WT levels (Fig. 6A-C).

**Figure 6.**
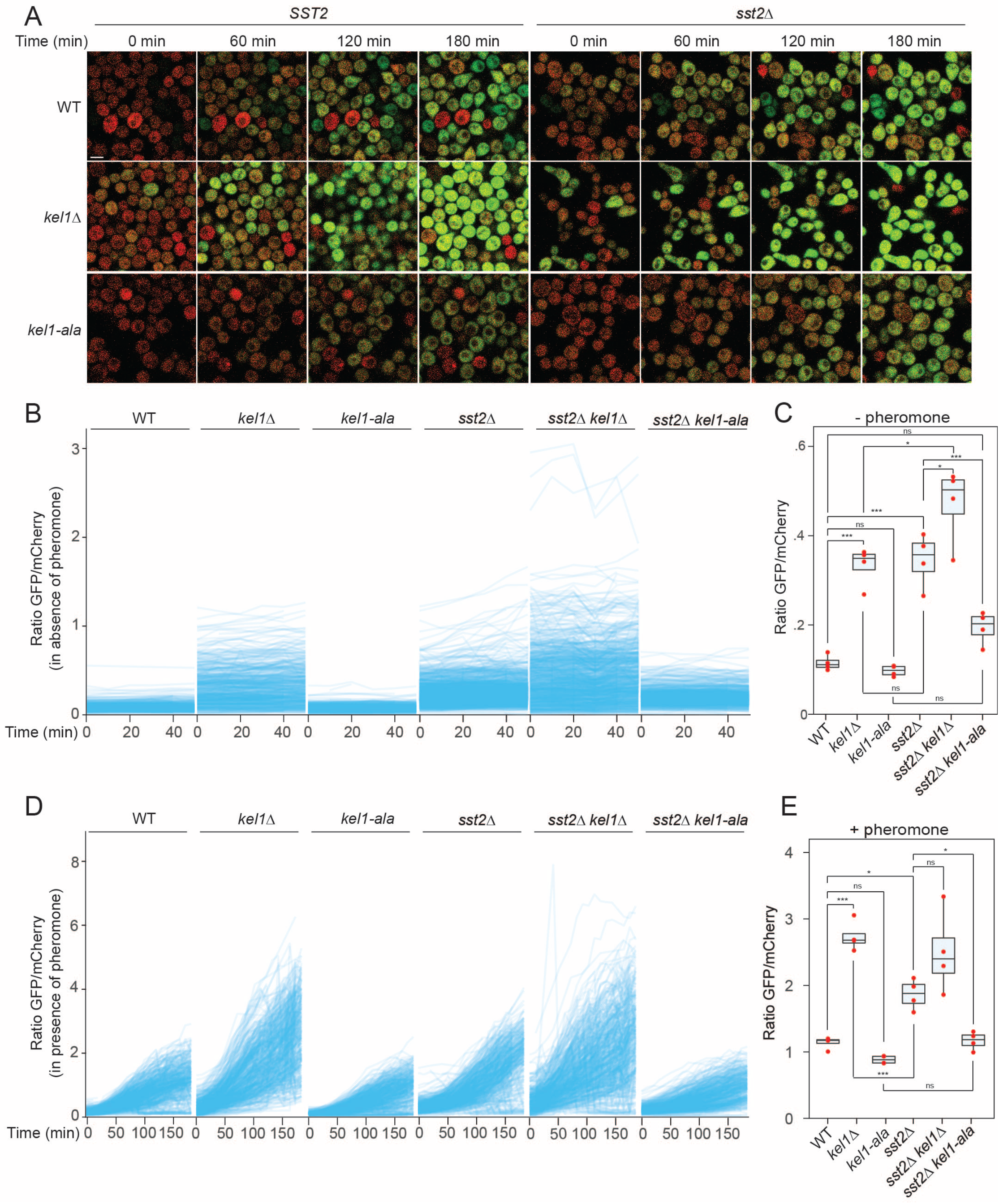
Kel1 dampens the pheromone signaling pathway. ***A,*** Representative micrographs of pheromone signaling time-courses. Populations of cells of thee indicated genotype carrying the reporter system *FUS1p*-GFP/*ADH1p*-mCherry were imaged by fluorescence microscopy at different timepoints in absence or presence of pheromone. Bar, 5 µM. ***B,*** Pheromone signal trajectories in untreated cells. Cells were imaged every 10 min. After image alignment, cells were identified, labeled and tracked over time (50 min). For each individual cell the ratio of GFP/mCherry intensities was measured and plotted (blue trajectories). The plot shows the cellular trajectories of one representative experiment. ***C,*** Median GFP/mCherry intensity in absence of pheromone. After 60 min in absence of pheromone, the median of the GFP/mCherry ratios was calculated for each strain and the values of 4 independent experiments (red dots) were plotted. Statistical significance was calculated using Tukey’s test on ANOVA fitted data, *, p<0.05; **, p<0.005; *** p<0.0001. ns, not significant. ***D,*** Pheromone signal trajectories in presence of pheromone (15 mg/l for 180 min). The experiment was performed in similar conditions as in (B). ***E,*** Median GFP/mCherry intensity in presence of pheromone (15 mg/l for 180 min). The analysis was performed as described in (C). *, p<0.05; **, p<0.005; *** p<0.0001. Four independent experiments were performed in which at least 31,000 cells were analyzed for each strain. Dead cells, which autofluoresce much more brightly than GFP-expressing living cells, were filtered out.

Next, we treated the different strains with pheromone and monitored cellular responses using single-cell time-lapse microscopy. Compared to the WT strain, the signal amplitude was higher in the majority of *kel1*Δ mutants (Fig. 6A,D). Quantification of the signal after 180 min of pheromone treatment supported these findings (Fig. 6E). Pheromone treatment resulted in a generally higher amplitude in *sst2*Δ cells than in WT cells (Fig. 6A-C), but strikingly, expression of *kel1-ala* in the *sst2*Δ background reduced the output of the pheromone signaling pathway to a level comparable to that of WT cells (Fig. 6A,D,E). These data are consistent with an inhibitory role for hypophosphorylated Kel1 downstream of Sst2.

### Kel1 suppresses noise in the pheromone signaling pathway

We noticed that there existed considerable cell-to-cell variability in the pheromone response of *kel1Δ* populations, suggesting that Kel1 may suppress noise in the pathway. We measured the overall signaling noise as the coefficient of variation (CV; i.e. the ratio between the standard deviation/mean) of the median population response before and after pheromone treatment. The dispersion (the extent to which a distribution is stretched or squeezed) in WT signaling was between 40-50% both before and after pheromone treatment, but increased slightly in the dynamic range of the pheromone response as previously reported (Fig. 7A,B) (Dixit et al., 2014). Interestingly, in untreated cell populations the signal dispersion was significantly higher in *kel1*Δ mutants than in WT cells, whereas there was no significant difference between *sst2*Δ and WT cells (Fig. 7A,B, see Fig. 7C for a dispersion time course), indicating that Kel1 is a suppressor of noise in the pheromone pathway. After 180 minutes of pheromone treatment we could no longer observe the increase in overall noise that we detected in populations of *kel1Δ* and *sst2*Δ mutants in absence of pheromone (Fig. 7B). This is likely a consequence of the elevated signaling and increase in average GFP expression in these mutants, which causes an expected decrease in CV for overdispersed distributions (Fig. 6)(Eling et al., 2019). The increased signaling variability in the *kel1Δ* mutant in the absence of pheromone led us to speculate whether Kel1 serves to suppress intrinsic signaling noise and spontaneous activation of the signaling pathway, while maintaining high signaling fidelity in response to pheromone. We used regression to estimate the intrinsic noise from non-reducible residual fluctuations over modeled signaling trajectories for individual cells (Hilfinger and Paulsson, 2011). By using a linear model in the linear range of the pheromone signaling response, we observed a significant increase in variation over the predicted signaling trajectories in the *kel1Δ* mutant and the *kel1Δ sst2Δ* double mutant compared to the WT strain (Fig. 7D,E), whereas the *sst2Δ* mutation alone did not significantly increase residual variation. Interestingly, expression of the *kel1-ala* allele strongly reduced variation to levels well below those of WT cells (Fig. 7D,E). Taken together, these results show that Kel1 suppresses noise, and it limits spontaneous signal transmission in the absence of pheromone.

**Figure 7.**
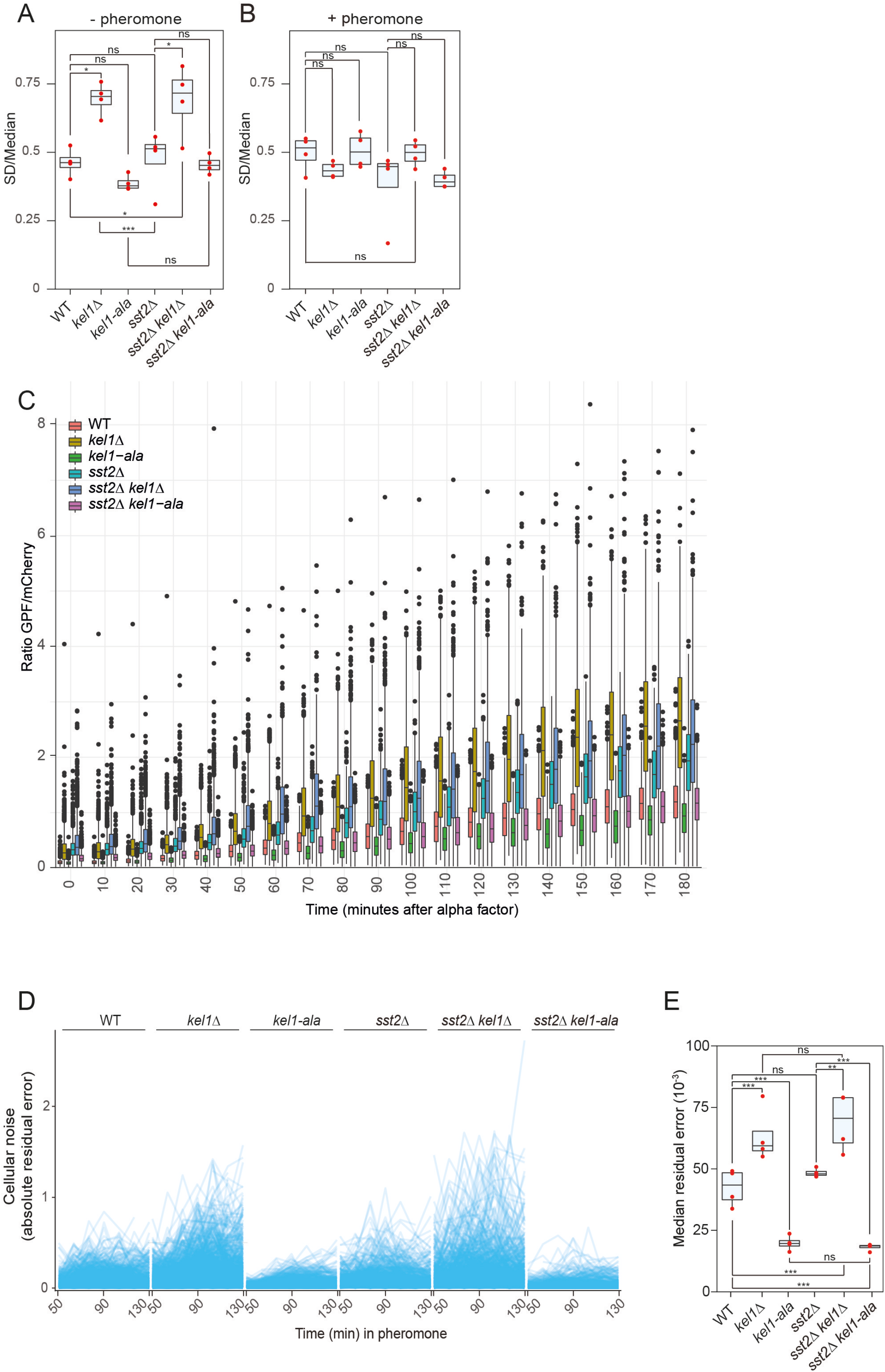
Kel1 prevents noise in the pheromone signaling pathway. ***A,*** Noise in the pheromone signaling pathway in absence of pheromone. Noise was computed as the ratio between the standard deviation and the median of the ratio of GFP and mCherry fluorescence intensities after 60 min in absence of pheromone from 4 independent experiments (red dots). Statistical significance was calculated using Tukey’s test on ANOVA fitted data, *, p<0.05; **, p<0.005; *** p<0.0001. ns, not significant. ***B,*** Noise in the pheromone signaling pathway in presence of pheromone (15 mg/l for 180 min), calculated as in (A). ***C,*** Signal dispersion in the pheromone signaling pathway upon pheromone treatment over time. ***D,*** Within-cell variations in the pheromone signaling pathway. Fluctuation trajectories of individual cells (blue lines) were calculated by computing the absolute error of the ratio of GFP and mCherry intensities from an averaged linear trajectory for each individual cell after addition of pheromone for 50-130 min. ***E,*** Median of the intrinsic residual error in the linear range (50-130 min) of the pheromone signaling pathway. The residual error was calculated as in (Fig. 7D) Statistical significance was calculated using Tukey’s test on ANOVA fitted data, *, p<0.05; **, p<0.005; *** p<0.0001. ns, not significant. Four independent experiments were performed in which at least 31,000 cells were analyzed for each strain. Autofluorescent dead cells were filtered out.

## Discussion

High levels of noise in signaling pathways can result in a substantial fitness cost, and therefore filtering mechanisms have evolved to suppress noise. Here, we show that hypophoshorylated Kel1 mediates noise suppression, inhibits spontaneous activation of the pathway, and prevents cell death during pheromone treatment. Fus3 is one putative Kel1 kinase, but other kinases also contribute. Several proline-directed kinases have been shown to have a role in the pheromone response, including the MAPKs Kss1 and Slt2 (Zarzov et al., 1996). The CDK Pho85, which appeared to be involved in Kel1 phosphorylation in vegetative cells, responds to certain environmental cues, such as nutrient status, and it will be interesting to study how Pho85 affects Kel1. Cdk1 is inhibited by pheromone treatment and therefore less likely to phosphorylate Kel1 (Peter and Herskowitz, 1994). Other CDKs include Bur1, Ctk1, Srb10 and Kin28, which are best known for transcription, although some also have non-transcriptional functions (Rother and Strasser, 2007; Zhu et al., 2016). Non-proline directed kinases that can target S/T-P sites include Tor1/2; the GSK3β homologs Mck1, Ygk3, Mrk1 and Rim11; and the LAMMER kinase Kns1. Thus, the number of possible permutations of combined kinase knockouts is large, which is also true for the number of potential combinations of alanine/aspartate substitutions. Systematic mapping of phosphosites and kinase mutagenesis will therefore be the focus of future studies. We expect this will provide insight into the upstream signals and pathways that converge upon Kel1 in parallel to the pheromone response pathway. It is possible that Kel1 integrates certain cell homeostasis signals, such as status of the cell wall, or nutrient availability, with polarized shmoo growth. Interestingly, it was recently shown that in response to mechanical stress Pkc1 prevents lysis of pheromone-treated cells by phosphorylating Ste5, resulting in dispersal of Ste5 from the site of polarized growth (van Drogen et al., 2019). We found that Kel1 also suppresses lysis during pheromone treatment and that it regulates Ste5 localization. Moreover, Kel1 was first identified in an overexpression screen for genes that overcome the fusion defect of cells expressing activated Pkc1 (Philips and Herskowitz, 1998). It will be interesting to investigate the relationship between Pkc1, Kel1 and Ste5 at the molecular level.

What could be the physiological relevance of Kel1 during mating? Under suboptimal mating conditions where mates are only transiently available, such as in dynamic aqueous environments, hypophoshorylated Kel1 may prevent unsuccessful mating attempts; unsuccessful mating is associated with a fitness cost (Banderas et al., 2016), Our working model is shown in Suppl. Figure S2A-D). We speculate that phosphorylation and thereby inhibition of Kel1 by Fus3 may help set up a feedback loop that boosts the pheromone response (Suppl. Fig. S2B); such feedback loops can act as bistable switches, resulting in rapid all-or-none changes in cellular states (Ferrell, 2002). Previous findings indicate Ste5-dependent switch-like behavior of the mating pathway (Malleshaiah et al., 2010; Paliwal et al., 2007); it will be interesting to test if and how Kel1 affects switch-like decision-making in the pheromone pathway.

Computer simulations and experimental studies have revealed mechanisms that provide robustness to the mating response (Chen et al., 2016; Howell et al., 2012), which is defined here as the persistence of a system’s behavior under conditions of uncertainty. Robustness is critical for efficient yeast mating and one way cells improve robustness is by reducing the sensitivity to pheromone, which is in part mediated by Sst2 (Chen et al., 2016). We found that the transcriptional response to pheromone in individual *kel1Δ* mutant cells fluctuated considerably over time, whereas *kel1-ala* mutants showed less fluctuations than WT cells. This suggests that hypophosphorylated Kel1 may provide robustness to the system. The phenotype of the *kel1Δ* mutant and the *kel1-ala* mutant masked the increase in signaling noise in *sst2Δ* mutants, indicating that proper control of Ste5 may be the dominant limiting factor for signal transmission fidelity and amplitude. We believe that the balance between phosphorylated and hypophosphorylated Kel1 is important for fine-tuning the output of the pathway by controlling Ste5.

### Limitations of the study

Although the phenotype of the *kel1-asp* allele resembled that of the *kel1Δ* mutation, it was generally less penetrant. This could be due to the fact that phosphomimicking residues do not always fully replicate the effect of phosphorylation (e.g. see (Paleologou et al., 2008)). For instance, substitution with Asp/Glu residues confers just a single negative charge, whereas phosphogroups confer multiple; there are differences in the distance of the negative charges to the protein backbone; and the pKa values are dissimilar. *kel1Δ* mutant cells may also have lost certain functions important for morphogenesis that are not regulated by phosphorylation, such as potential adapter functions conferred by the Kelch propeller and the coiled coil domains. Other limitations of our study that remain to be addressed are to identify the phosphatase that targets Kel1 and to determine how phosphorylation regulates Kel1 activity at the molecular level, although it does not affect the expression or localization of Kel1 (Suppl. Fig. S1H,I). The mechanism by which Kel1 controls activity of the mating pathway also remains to be established. Kel1 consists of a Kelch propeller and coiled coil domains, which are known to mediate multimerization and protein-protein interactions. Given that Kel1 can physically interact with Ste2, Sst2, Far1, Ste5, Fus3, we speculate that Kel1 may form a phospho-dependent platform that integrates signals to regulate the pheromone pathway. Future studies will focus on unraveling the molecular mechanism by which Kel1 regulates membrane recruitment of Ste5 and Far1.

In conclusion, we have shown that Kel1 is an important regulator of the mating pathway with a major function in noise suppression.

## Acknowledgments

Fus3 plasmids were kindly provided by Dr. Dohlman (University of North Carolina, Chapel Hill, USA). This work was supported by grants from the Norwegian Cancer Society (project numbers 182524 and 208012), the Norwegian Health Authority South-East (2017064, 2017072, 2018012, 2019096) and the Norwegian Research Council (261936, 301268). This work was partly supported by the Research Council of Norway through its Centres of Excellence funding scheme, project number 262652.

## Author contributions

Conceptualization: JME, IG. Methodology: IG, LRA, PC, SMO and JME. Investigation: IG, PC, SMO, JE, MPC. Formal analysis: IG, LRA, JME, ANA, PC, SMO, JE. Writing original draft: IG, ANA, JME. Writing – Review & Editing: IG, ANA, PC, SMO, JME. Visualization: IG, PC, SMO, LRA, JE. Supervision: IG, JME, SOB, PC. Project Administration: IG, JME, PC. Funding Acquisition: JME, IG, PC.

## Declaration of Interests

The authors declare no competing interests.

## STAR Methods

### RESOURCE AVAILABILITY

#### Lead contact

Further information and requests for resources and reagents should be directed to and will be fulfilled by the lead contact, Jorrit M. Enserink.

#### Materials availability

Plasmids generated in this study will be shared by the lead contact upon request.

#### Data and code availability

Mass-spectrometry data are deposited to the ProteomeXchange Consortium via PRIDE repository with accession number PXD020833.

All data reported in this paper will be shared by the lead contact upon request.

Any additional information required to reanalyze the data reported in this paper is available from the lead contact upon request.

### EXPERIMENTAL MODEL AND SUBJECT DETAILS

*S. cerevisiae* strains were grown in at 30°C until mid-log phase in standard yeast extract peptone dextrose (YPD) medium or in synthetic medium supplemented with relevant amino acids. Strains were derived directly from the S288c strains BY4741 (Brachmann et al., 1998) and RDKY3032 (Flores-Rozas and Kolodner, 1998) using either standard gene-replacement methods or intercrossing. See Suppl. Table S1 for strains.

### METHOD DETAILS

#### Strain construction

To construct *kel1-ala* and *kel1-asp* mutants, *KEL1* was first replaced with the *URA3* gene. Next, the *URA3* on the strain *kel1::URA3* was replaced by *kel1-ala* and *kel1-asp* alleles, which were PCR-amplified from plasmids containing *in vitro* synthesized *kel1* alleles (GenScript) in which the S/T-P sites were replaced with either alanine or aspartate residues, respectively. In some cases where two or three serine/threonine residues preceded a proline, all were substituted. Note that *kel1-asp* differs slightly from *kel1-ala* in terms of mutated sites (additional substitution of T65 and S688). Alanine substitutions in Kel1-ala: T18, S34, T35, T47, S66, S67, S121, T178, T293, T465, T467, T477, T480, S503, T549, T559, T604, S613, T626, S689, S748. Aspartate substitutions in Kel1-asp: T18, S34, T35, T47, T65, S66, S67, S121, T178, T293, T465, T467, T477, T480, S503, T549, T559, T604, S613, T626, S688, S689, S748.

#### Plasmid construction

The pFA6-mNG plasmid was generated by replacing the eGFP coding sequence from pYM-28 (Janke et al., 2004) by codon-optimized monomeric NeonGreen (mNG) DNA sequence (Shaner et al., 2013) synthesized by GenScript. mNG sequence was amplified using primers NG-FW and NG-RV (for primer sequences see Key Resources Table), which also include *HindIII* and *BamHI* sites for the tag replacement. pGEX-5X-1 plasmids containing Kel1-NT, Kel1-NT-ala, Kel1-mid and Kel1-mid-ala were synthesized by GenScript. Plasmids pGEX-2T-GST-Fus3 and pGEX-2T-GST-Fus3-KD for purification of Fus3 and kinase-dead Fus3-KD have been previously described (Parnell et al., 2005).

#### Pheromone treatment

Cells were treated with 15mg/L of alpha factor (custom synthesized by GenScript) for 2 hrs unless otherwise indicated.

#### Mating efficiency assay

To evaluate mating efficiency, 1ml of mid-log-phase MATa and MATalpha cells carrying complementary markers were mixed and incubated at 30°C in absence of agitation for 4 hours. 100μl of five consecutive serial dilutions (1:10) of the crosses were plated on YPD plates and diploid selective media (YPD supplemented with G418 200μg/ml (Sigma-Aldrich) and nourseothricin 100μg/ml (WERNER BioAgents GmbH)). After 2 days of incubation at 30°C the colonies of the plates were counted using the Colony Counter mobile application (Promega). See Suppl. Table S1 for strains.

#### FITC staining

Cells were harvested, washed with PBS and stained for 10 min in the dark with 1mg/ml FITC (Sigma-Aldrich) in carbonate buffer (0.1M, pH 9.5). Cells were washed 3 times with PBS and visualized by fluorescence microscopy as described below.

#### Flow cytometry

GFP and mCherry-expressing cells were harvested and fixed with 4% paraformaldehyde for 15 min at room temperature. Paraformaldehyde was quenched with glycine 0.5M for 15 min at room temperature. Cells were washed 2 times with PBS, resuspended in PBS and sonicated for 10 seconds at 30% amplitude (Hielscher Ultrasonics UP 400S). Flow cytometry was performed using an LSR II Flow Cytometer (BD Biosciences). Data analysis and plotting was performed in R.

#### Mass spectrometry

Cells untreated or treated with of alpha-factor (15 mg/L) for 2 hrs. were collected, washed with cold PBS and lysed in lysis buffer (100mM Tris-HCl pH8, 150mM NaCl, 1mM EDTA, 1% Triton X-100, 10% Glycerol, 2mM DTT, protease and phosphatase inhibitor cocktails). After centrifugation, supernatants were immunoprecipitated by mixing 800 µg of cell extract with 20 µL of anti-Myc-Tag magnetic bead (Cell Signaling). After 2 hrs. incubation at 4 °C, beads were washed once with wash buffer A (100mM Tris-HCl pH8, 150mM NaCl, 1mM EDTA, 5% Glycerol, 2mM DTT, protease and phosphatase inhibitor cocktails), twice with wash buffer B (100mM Tris-HCl pH8, 150mM NaCl, 1mM EDTA, 2mM DTT, protease and phosphatase inhibitor cocktails) and twice with PBS. The proteins were digested directly on-beads with trypsin, and the resulting peptides were purified with C18-microcolumns. For mass spectrometry analysis 3µl of each sample was injected in triplicates to nLC-MS/MS (nEASY LC-QExactive Plus) using a 50cm C18 nLC column and 60 min separation gradient.

#### Cell lysis, immunoprecipitation, λ phosphatase treatment and western blotting

Immunoprecipitation and western blotting were performed as previously described (Chymkowitch et al., 2012; Chymkowitch et al., 2017; Herrera et al., 2018). For coimmunoprecipitations, cells were washed with ice-cold PBS, resuspended in lysis buffer (100mM Tris-HCl pH8, 150mM NaCl, 1mM EDTA, 1% Triton X-100, 10% Glycerol, 2mM DTT, protease and phosphatase inhibitor cocktails) and lysed by vortexing with glass beads followed by centrifugation (14,000 rpm at 4°C for 10 min) to remove cell debris. Equal amounts of proteins were used for immunoprecipitation using magnetic beads conjugated covalently to relevant antibodies. After extensive washing with lysis buffer, coimmunoprecipitated proteins were resolved by SDS-PAGE and analyzed by western blotting with the indicated antibodies. For λ phosphatase treatments, cells were harvested and lysed as described above, magnetic beads were washed 2 times with lysis buffer without phosphatase inhibitors. Samples were treated with 800 units of λ-phosphatase (New England Biolabs) for 30 min after which the enzyme was heat-inactivated (5 min at 95°C) and the proteins resolved by Phos-tag SDS-PAGE (see below) and analyzed by western blotting.

#### Phos-tag gel electrophoresis

Proteins from synchronized cells samples were resuspended in ice-cold RIPA Buffer (50 mM Tris HCl, 150 mM NaCl, 1.0% (v/v) NP-40, 0.5% (w/v) Sodium Deoxycholate, 1.0 mM EDTA, 0.1% (w/v) SDS and 0.01% (w/v) sodium azide at a pH of 7.4), lysed in a bead mill and centrifuged (14,000 rpm at 4°C for 10 min) before immunoprecipitation using magnetic beads conjugated covalently to relevant antibodies as previously described (Herrera et al., 2018). Equal amounts of sample were loaded onto 5% acrylamide (BioRad) containing 6-15 µM Phos-tag (Wako chemicals) gels, followed by electrophoresis for 12 hours at 50 V. Gels were washed three times for 10 min with transfer buffer containing 5 mM EDTA and three times for 10 min with transfer buffer without containing EDTA. The gel was transferred to a membrane and protein phosphorylation was analyzed by western blotting as described above. Protein densitometric quantifications were performed using FIJI software (Schindelin et al., 2012) and analyzed using R.

#### Purification of Recombinant Proteins

GST alone, GST-Kel1-NT, GST-Kel1-NT-Ala, GST-Kel1-Mid, GST-Kel1-Mid-Ala GST-Fus3 and GST-Fus3-KD (Parnell et al., 2005) were expressed 2.5 hours at 37°C in *E. coli* BL21 bacteria induced with 400 μM isopropyl-β-D-thio-galactoside (IPTG). Cells were harvested and washed with PBS before sonication in ice-cold lysis buffer [PBS, 1% (vol/vol) Triton X-100, 2 mM DTT, and protease inhibitor mixture]. The lysate was centrifuged, and the supernatant was incubated for 1h at 4 °C with glutathione Sepharose 4B beads (Amersham). After three washes with PBS, GST fusion proteins were collected in elution buffer (50 mM Tris pH 8.0, 10 mM reduced glutathione, 2 mM DTT and protease inhibitor mixture and stored at −80°C.

#### In Vitro Kinase Assays

Kinase assays were performed as previously described (Chymkowitch et al., 2012). Briefly, two micrograms of purified recombinant substrate protein was incubated with [γ-32P] ATP (0.14 mM) in the presence of 300 ng of purified recombinant GST-Fus3 or GST-Fus3-KD (Parnell et al., 2005) in kinase buffer [20 mM Hepes-Na+ (pH 7.9), 20 mM Tris·HCl (pH 7.9), 5 mM MgCl2, 30 mM KCl, and 4% (vol/vol) glycerol]. After 40 min at 30°C, reactions were stopped with Laemmli sample buffer. Samples were boiled and resolved by SDS-PAGE and stained with Coomassie Brilliant Blue. Radiolabelled GST fusions were analyzed using a phosphorimager (Amersham Typhoon Biomolecular Imager from GE Healthcare Life Sciences).

#### Microscopy

DIC images were acquired on a Leica DM600B microscope using a 63X/1.40NA immersion objective. The microscope was connected to a Hamamatsu C9100-14 camera controlled by Leica Application Suite software.

FITC staining Far1-mNG and brightfield images were acquired using a Zeiss AXIO Scope.A1 microscope containing a 63X/1.40NA immersion objective and a Leica DFC camera. The microscope was controlled by Micro-Manager software (Edelstein et al., 2010). Ste5-mNG images were obtained using a Zeiss AXIO Observer.Z1 microscope containing a 63X/1.40NA immersion objective. The microscope was equipped with a Hamamatsu ORCA-Flash4.0 camera and temperature control and it was controlled by Micro-Manager software.

Brightfield and Airyscan confocal images were obtained using a Zeiss LSM880 Airyscan microscope (Carl Zeiss MicroImaging GmbH, Jena, Germany) using a 63X oil DICII objective, and the Airyscan detector in super resolution mode. Images were deconvolved using ZEISS ZEN Software.

#### Synthetic genetic array

The SGA query strain Y8205 (*can1::STE2pr-Sp_his5 lyp1::STE3pr-LEU2 his3Δ1 leu2Δ0 ura3Δ0*; kind gift from C. Boone, University of Toronto, Canada) either harboring the *kel1Δ* or the *kel1-ala* mutation was crossed with a collection of deletions of non-essential genes (YKO collection) and with a collection of mutants with reduced mRNA levels of essential yeast genes (DAmP collection) according to (Tong et al., 2001). Briefly, mutations of interest were linked to the natNT2 selection marker, while mutations in the collections were selectable using the kanMX antibiotic resistance cassette. After mating and sporulation, the spores were transferred to medium that enables growth of MATa meiotic progeny. In the final step, double mutants were obtained after selection on medium containing kanamycin and nourseothricin. Each cross was done in quadruplicate on 1536-format plates. All double mutants were grown at 30°C and 37°C and imaged after 2 days. Image analysis and scoring were done with SGAtools (Wagih et al., 2013).

The phenotype enrichment score (Fig. 3B) was computed in R using the bc3net library (de Matos Simoes et al., 2012) and it was calculated over genes annotated with phenotypes in invasive growth, response to pheromone, pheromone sensitivity, mating projection morphology, cell shape, endomembrane system morphology, cell wall morphology and cytoskeleton morphology. All annotations were downloaded from the Yeast Phenotype Ontology database (https://www.yeastgenome.org/ontology/phenotype/ypo).

#### RT-qPCR

RT-qPCR experiments were performed as previously described (Chymkowitch et al., 2012). For primer sequences see Key Resources Table. Briefly, for the RNA extraction the RNeasy Mini Kit (QIAGEN) was used the mechanical disruption of cell pellets as indicated in the manufacturer’s protocol. For cDNA synthesis the QuantiTect Reverse Transcription Kit (QIAGEN) was used. RT-qPCR was performed using the StepOnePlus™ Real-Time PCR System (Applied Biosystems™ by Thermo Fisher Scientific, Cat. No. 4376600) and Power SYBR™ Green PCR Master Mix (Applied Biosystems™ by Thermo Fisher Scientific).

### QUANTIFICATION AND STATISTICAL ANALYSIS

#### Hypothesis testing

Two-proportion z-test was used to calculate the statistical significance of assays in which proportions of cells from different observations and independent experiments were measured (such as viability assays, shmoo morphology and mating efficiency).

To estimate the dispersion of the population, a fraction of 100 observations were sampled 1000 times from the total and the standard deviations were calculated on the bootstrapped sample. Two-proportion z-tests and standard deviations were calculated in R from at least 3 independent experiments containing at least 100 observations.

t-test was used to calculate the statistical significance of assays that generate one single value per observation (such as ellipticity measurements, mRNA levels and fluorescence quantification). t-tests were calculated in R from at least 3 independent experiments.

ANOVA and Tukey’s tests were used to calculate the statistical significance of the time-course assays. ANOVA and Tukey’s tests were calculated in R from 4 independent experiments in which at least 31000 cells were analyzed for each strain.

Significance was classified in four different categories according to the p-value (unless stated otherwise in the figure legend):

- Highly significant: p-value < 0.001 (***)
- Very significant: 0.01 > p-value > 0.001 (**)
- Significant: 0.05 > p-value > 0.01 (*)
- Non significant: p-value > 0.05 (ns)

### Supplemental Table Titles and Legends

**Supplemental Dataset S1.** CSV file containing data from the SGA screen with *kel1Δ* and *kel1-ala* query alleles. Related to Figure 3A.

**Supplemental Dataset S2.** CSV file containing data from the MS proteomics experiments to analyze the Kel1 interactome. Related to Figure 4A.

---

**Figure S1.**
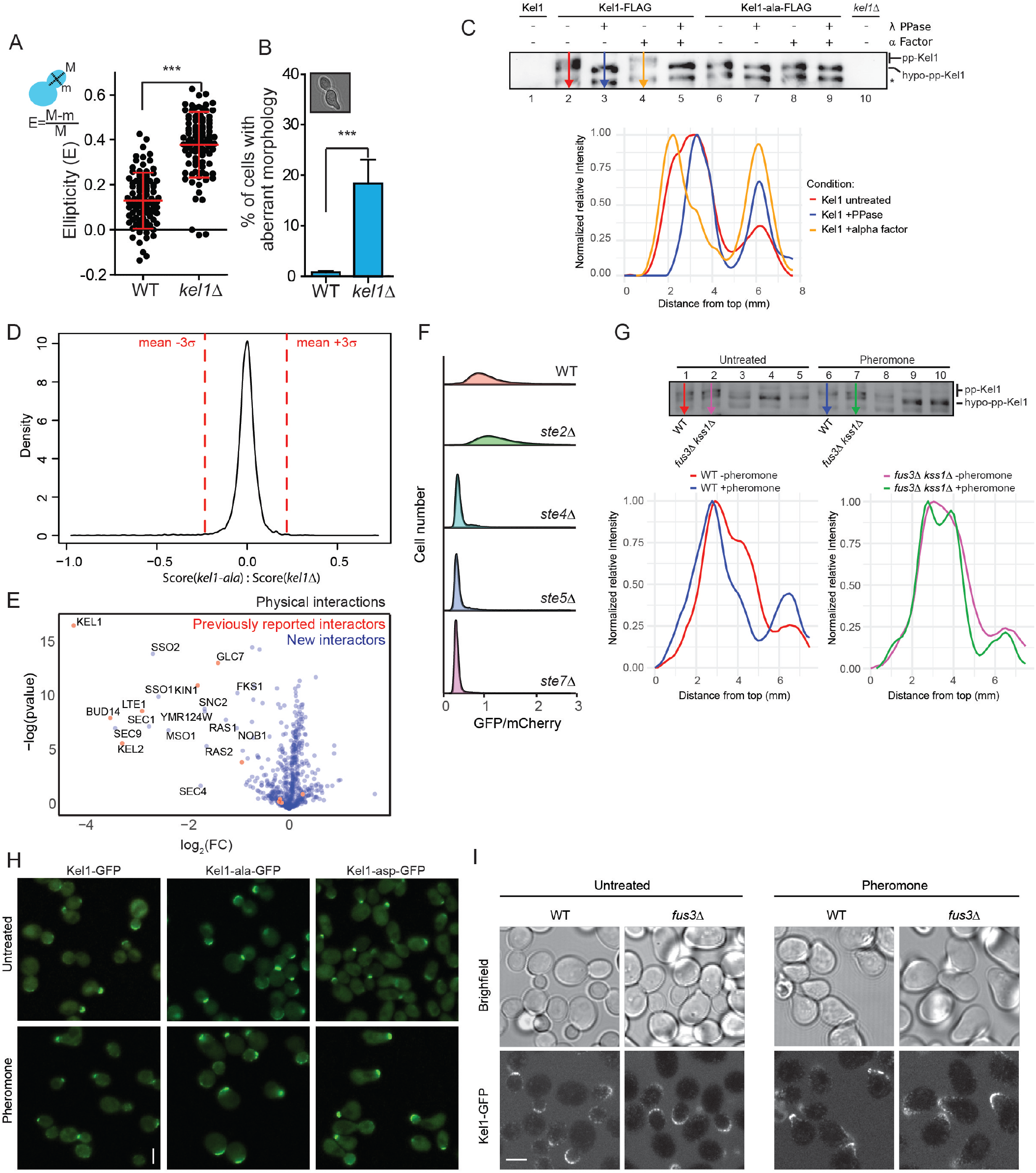
***A,*** Kel1 suppresses elongated bud growth. WT cells and *kel1*Δ mutants were grown to log phase, imaged by light microscopy and bud ellipticity was calculated as indicated in the figure. Related to Figure 1. ***B*,** Loss of *KEL1* results in morphological aberrancies. Quantification of morphology of cells imaged in S1A. Related to Figure 1. ***C*,** Densitometric quantification of Kel1 phosphorylation of the WB shown in Figure 2B. Pixel intensity across the lanes in the WB (WB copied from Fig. 2B) was quantified as indicated with the arrows, followed by normalization of the data between 0 and 1, where 1 is the most intense pixel on the lane. Plots were aligned to maximum intensity peak of the lower band, the Mw of which does not change in response to treatment. ***D,*** Statistical cut-off of mutants analyzed in Figure 3A. ***E,*** Overview of physical interactions with Kel1 (see Figure 4A for description of the experiment). ***F,*** Noise in the pheromone signaling pathway is generated downstream of Ste2. The ratio of GFP/mCherry in the indicated mutants was determined by fluorescence microscopy. Related to Figure 6. ***G***, Densitometric quantification of Kel1 phosphorylation of the WB shown in Figure 4C. Pixel intensity across the lanes in the WB (copied from Fig. 4C) was quantified as in S1C. ***H,*** Expression and localization of Kel1 are not affected by phosphorylation. Cells were grown to log phase before treatment with pheromone (15mg/l) for 2 hrs, after cells were imaged by fluorescence microscopy. Bar, 5µm. Related to Figure 2 and 3. ***I***, Kel1 localizes to sites of polarized growth in *fus3*Δ mutants. Cells were treated and imaged as in (H). Bar, 5µm. Related to Figure 2 and 3.

**Supplemental Figure S2.**
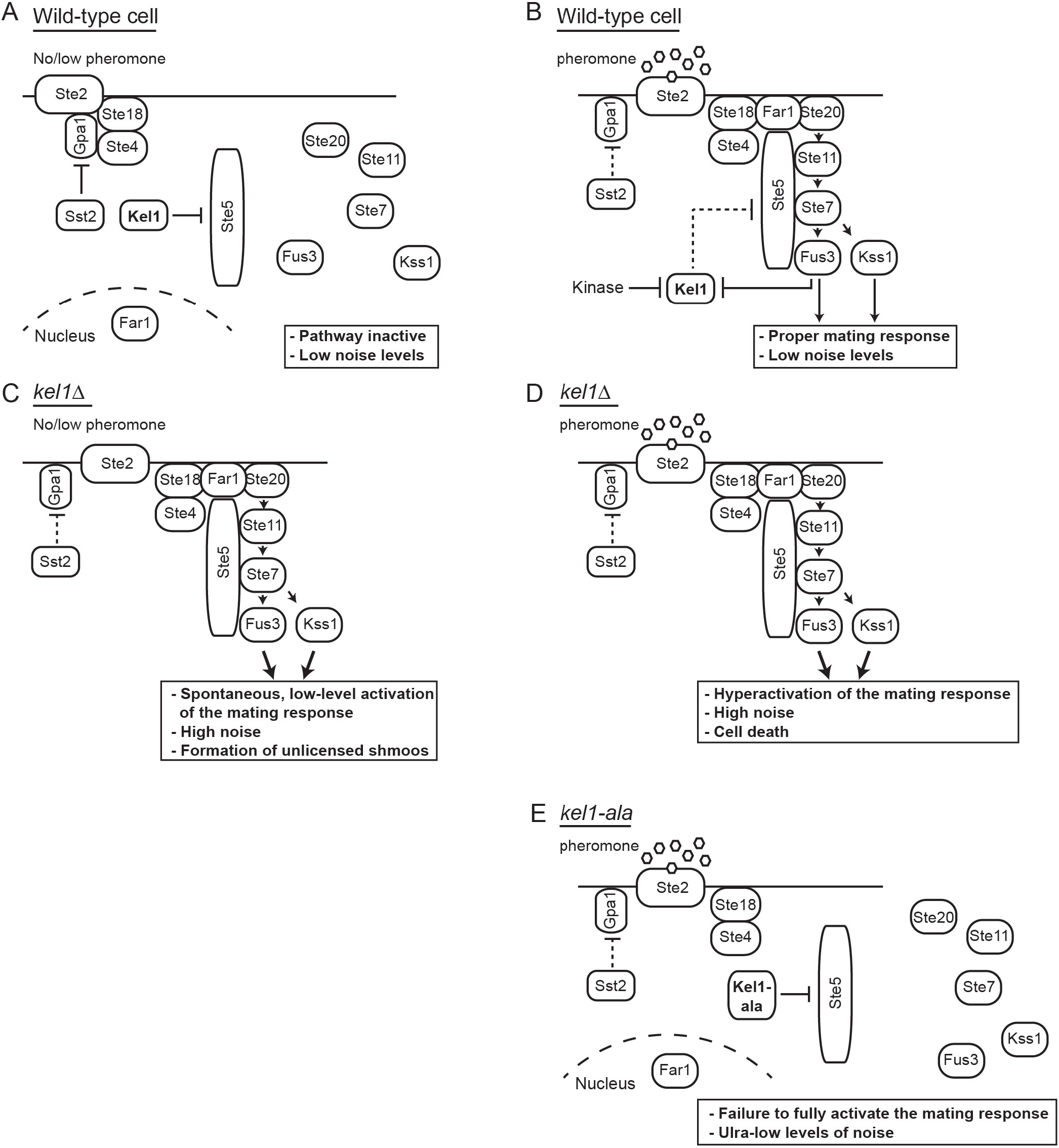
Model for regulation of the mating response by Kel1 phosphorylation (related to Figures 1-7 in the paper). ***A,*** In absence of pheromone, hypophosphorylated Kel1 prevents spontaneous activation of the pheromone response by inhibiting recruitment of Ste5 and Far1 to the membrane, whereas Sst2 prevents spontaneous activation of the pathway by inhibiting Gpa1. ***B,*** Pheromone treatment results in dissociation of the beta-gamma subunit of the G protein, which induces membrane recruitment of Ste5 and Far1. This results in activation of Fus3, which phosphorylates and thereby inactivate Kel1, allowing for recruitment of additional Ste5 to induce robust activation of the pathway. Other kinases are also important, which remain to be identified. ***C,*** In *kel1*Δ cells, Ste5 and Far localize spontaneously to the membrane even in absence of pheromone, resulting in increased transcription of pheromone response genes, shmoo formation and noise in the pathway. ***D,*** Pheromone treatment of cells lacking Kel1 results in hyperactivation of the mating pathway, high levels of noise, and cell death. ***E,*** Expression of non-phosphorylatable Kel1 inhibits membrane recruitment of Ste5 and Far1, thereby suppressing the pheromone pathway and strongly reducing noise.

## Notes

### Competing Interest Statement

The authors have declared no competing interest.

## References

Alvaro, C.G., and Thorner, J. (2016). Heterotrimeric G Protein-coupled Receptor Signaling in Yeast Mating Pheromone Response. J Biol Chem 291, 7788–7795.

Apanovitch, D.M., Iiri, T., Karasawa, T., Bourne, H.R., and Dohlman, H.G. (1998). Second site suppressor mutations of a GTPase-deficient G-protein alpha-subunit -Selective inhibition of G beta gamma-mediated signaling. Journal of Biological Chemistry 273, 28597–28602.

Balazsi, G., van Oudenaarden, A., and Collins, J.J. (2011). Cellular decision making and biological noise: from microbes to mammals. Cell 144, 910–925.

Banderas, A., Koltai, M., Anders, A., and Sourjik, V. (2016). Sensory input attenuation allows predictive sexual response in yeast. Nature Communications 7, 12590.

Baryshnikova, A., Costanzo, M., Dixon, S., Vizeacoumar, F.J., Myers, C.L., Andrews, B., and Boone, C. (2010). Synthetic genetic array (SGA) analysis in Saccharomyces cerevisiae and Schizosaccharomyces pombe. Methods Enzymol 470, 145–179.

Brachmann, C.B., Davies, A., Cost, G.J., Caputo, E., Li, J., Hieter, P., and Boeke, J.D. (1998). Designer deletion strains derived from Saccharomyces cerevisiae S288C: a useful set of strains and plasmids for PCR-mediated gene disruption and other applications. Yeast 14, 115–132.

Breslow, D.K., Cameron, D.M., Collins, S.R., Schuldiner, M., Stewart-Ornstein, J., Newman, H.W., Braun, S., Madhani, H.D., Krogan, N.J., and Weissman, J.S. (2008). A comprehensive strategy enabling high-resolution functional analysis of the yeast genome. Nat Methods 5, 711–718.

Butty, A.C., Pryciak, P.M., Huang, L.S., Herskowitz, I., and Peter, M. (1998). The role of Far1p in linking the heterotrimeric G protein to polarity establishment proteins during yeast mating. Science 282, 1511–1516.

Chen, W., Nie, Q., Yi, T.M., and Chou, C.S. (2016). Modelling of Yeast Mating Reveals Robustness Strategies for Cell-Cell Interactions. PLoS Comput Biol 12, e1004988.

Choudhury, S., Baradaran-Mashinchi, P., and Torres, M.P. (2018). Negative Feedback Phosphorylation of Ggamma Subunit Ste18 and the Ste5 Scaffold Synergistically Regulates MAPK Activation in Yeast. Cell Rep 23, 1504–1515.

Chymkowitch, P., Eldholm, V., Lorenz, S., Zimmermann, C., Lindvall, J.M., Bjoras, M., Meza-Zepeda, L.A., and Enserink, J.M. (2012). Cdc28 kinase activity regulates the basal transcription machinery at a subset of genes. Proc Natl Acad Sci U S A 109, 10450–10455.

Chymkowitch, P., Nguea, P.A., Aanes, H., Robertson, J., Klungland, A., and Enserink, J.M. (2017). TORC1-dependent sumoylation of Rpc82 promotes RNA polymerase III assembly and activity. Proc Natl Acad Sci U S A 114, 1039–1044.

Colman-Lerner, A., Gordon, A., Serra, E., Chin, T., Resnekov, O., Endy, D., Pesce, C.G., and Brent, R. (2005). Regulated cell-to-cell variation in a cell-fate decision system. Nature 437, 699–706.

de Matos Simoes, R., Tripathi, S., and Emmert-Streib, F. (2012). Organizational structure and the periphery of the gene regulatory network in B-cell lymphoma. BMC Syst Biol 6, 38.

Dixit, G., Kelley, J.B., Houser, J.R., Elston, T.C., and Dohlman, H.G. (2014). Cellular noise suppression by the regulator of G protein signaling Sst2. Mol Cell 55, 85–96.

Dohlman, H.G., Apaniesk, D., Chen, Y., Song, J.P., and Nusskern, D. (1995). Inhibition of G-Protein Signaling by Dominant Gain-of-Function Mutations in Sst2p, a Pheromone Desensitization Factor in Saccharomyces-Cerevisiae. Molecular and Cellular Biology 15, 3635–3643.

Edelstein, A., Amodaj, N., Hoover, K., Vale, R., and Stuurman, N. (2010). Computer control of microscopes using microManager. Curr Protoc Mol Biol Chapter 14, Unit14 20.

Eling, N., Morgan, M.D., and Marioni, J.C. (2019). Challenges in measuring and understanding biological noise. Nat Rev Genet 20, 536–548.

Elion, E.A., Satterberg, B., and Kranz, J.E. (1993). FUS3 phosphorylates multiple components of the mating signal transduction cascade: evidence for STE12 and FAR1. Mol Biol Cell 4, 495–510.

Ferrell, J.E., Jr. (2002). Self-perpetuating states in signal transduction: positive feedback, double-negative feedback and bistability. Curr Opin Cell Biol 14, 140–148.

Flores-Rozas, H., and Kolodner, R.D. (1998). The Saccharomyces cerevisiae MLH3 gene functions in MSH3-dependent suppression of frameshift mutations. Proc Natl Acad Sci U S A 95, 12404–12409.

Giaever, G., Chu, A.M., Ni, L., Connelly, C., Riles, L., Veronneau, S., Dow, S., Lucau-Danila, A., Anderson, K., Andre, B., et al. (2002). Functional profiling of the Saccharomyces cerevisiae genome. Nature 418, 387–391.

Gould, C.J., Chesarone-Cataldo, M., Alioto, S.L., Salin, B., Sagot, I., and Goode, B.L. (2014). Saccharomyces cerevisiae Kelch proteins and Bud14 protein form a stable 520-kDa formin regulatory complex that controls actin cable assembly and cell morphogenesis. J Biol Chem 289, 18290–18301.

Herrera, M.C., Chymkowitch, P., Robertson, J.M., Eriksson, J., Boe, S.O., Alseth, I., and Enserink, J.M. (2018). Cdk1 gates cell cycle-dependent tRNA synthesis by regulating RNA polymerase III activity. Nucleic Acids Res 46, 11698–11711.

Hilfinger, A., and Paulsson, J. (2011). Separating intrinsic from extrinsic fluctuations in dynamic biological systems. Proceedings of the National Academy of Sciences 108, 12167–12172.

Howell, A.S., Jin, M., Wu, C.F., Zyla, T.R., Elston, T.C., and Lew, D.J. (2012). Negative feedback enhances robustness in the yeast polarity establishment circuit. Cell 149, 322–333.

Janke, C., Magiera, M.M., Rathfelder, N., Taxis, C., Reber, S., Maekawa, H., Moreno-Borchart, A., Doenges, G., Schwob, E., Schiebel, E., et al. (2004). A versatile toolbox for PCR-based tagging of yeast genes: new fluorescent proteins, more markers and promoter substitution cassettes. Yeast 21, 947–962.

Kaern, M., Elston, T.C., Blake, W.J., and Collins, J.J. (2005). Stochasticity in gene expression: from theories to phenotypes. Nat Rev Genet 6, 451–464.

Lehner, B. (2008). Selection to minimise noise in living systems and its implications for the evolution of gene expression. Mol Syst Biol 4, 170.

Li, X., Gerber, S.A., Rudner, A.D., Beausoleil, S.A., Haas, W., Villen, J., Elias, J.E., and Gygi, S.P. (2007). Large-scale phosphorylation analysis of alpha-factor-arrested Saccharomyces cerevisiae. Journal of proteome research 6, 1190–1197.

Malleshaiah, M.K., Shahrezaei, V., Swain, P.S., and Michnick, S.W. (2010). The scaffold protein Ste5 directly controls a switch-like mating decision in yeast. Nature 465, 101–105.

Metzger, B.P., Yuan, D.C., Gruber, J.D., Duveau, F., and Wittkopp, P.J. (2015). Selection on noise constrains variation in a eukaryotic promoter. Nature 521, 344–347.

Nern, A., and Arkowitz, R.A. (1999). A Cdc24p-Far1p-Gbetagamma protein complex required for yeast orientation during mating. J Cell Biol 144, 1187–1202.

Oehlen, L.J., McKinney, J.D., and Cross, F.R. (1996). Ste12 and Mcm1 regulate cell cycle-dependent transcription of FAR1. Mol Cell Biol 16, 2830–2837.

Paleologou, K.E., Schmid, A.W., Rospigliosi, C.C., Kim, H.Y., Lamberto, G.R., Fredenburg, R.A., Lansbury, P.T., Jr., Fernandez, C.O., Eliezer, D., Zweckstetter, M., et al. (2008). Phosphorylation at Ser-129 but not the phosphomimics S129E/D inhibits the fibrillation of alpha-synuclein. J Biol Chem 283, 16895–16905.

Paliwal, S., Iglesias, P.A., Campbell, K., Hilioti, Z., Groisman, A., and Levchenko, A. (2007). MAPK-mediated bimodal gene expression and adaptive gradient sensing in yeast. Nature 446, 46–51.

Parnell, S.C., Marotti, L.A., Jr., Kiang, L., Torres, M.P., Borchers, C.H., and Dohlman, H.G. (2005). Phosphorylation of the RGS protein Sst2 by the MAP kinase Fus3 and use of Sst2 as a model to analyze determinants of substrate sequence specificity. Biochemistry 44, 8159–8166.

Peter, M., and Herskowitz, I. (1994). Direct inhibition of the yeast cyclin-dependent kinase Cdc28-Cln by Far1. Science 265, 1228–1231.

Philips, J., and Herskowitz, I. (1998). Identification of Kel1p, a kelch domain-containing protein involved in cell fusion and morphology in Saccharomyces cerevisiae. J Cell Biol 143, 375–389.

Raser, J.M., and O’Shea, E.K. (2005). Noise in gene expression: origins, consequences, and control. Science 309, 2010–2013.

Repetto, M.V., Winters, M.J., Bush, A., Reiter, W., Hollenstein, D.M., Ammerer, G., Pryciak, P.M., and Colman-Lerner, A. (2018). CDK and MAPK Synergistically Regulate Signaling Dynamics via a Shared Multi-site Phosphorylation Region on the Scaffold Protein Ste5. Mol Cell 69, 938–952 e936.

Roberts, C.J., Nelson, B., Marton, M.J., Stoughton, R., Meyer, M.R., Bennett, H.A., He, Y.D., Dai, H., Walker, W.L., Hughes, T.R., et al. (2000). Signaling and circuitry of multiple MAPK pathways revealed by a matrix of global gene expression profiles. Science 287, 873–880.

Rother, S., and Strasser, K. (2007). The RNA polymerase II CTD kinase Ctk1 functions in translation elongation. Genes Dev 21, 1409–1421.

Schindelin, J., Arganda-Carreras, I., Frise, E., Kaynig, V., Longair, M., Pietzsch, T., Preibisch, S., Rueden, C., Saalfeld, S., Schmid, B., et al. (2012). Fiji: an open-source platform for biological-image analysis. Nat Methods 9, 676–682.

Severin, F.F., and Hyman, A.A. (2002). Pheromone induces programmed cell death in S. cerevisiae. Curr Biol 12, R233–235.

Shaner, N.C., Lambert, G.G., Chammas, A., Ni, Y., Cranfill, P.J., Baird, M.A., Sell, B.R., Allen, J.R., Day, R.N., Israelsson, M., et al. (2013). A bright monomeric green fluorescent protein derived from Branchiostoma lanceolatum. Nat Methods 10, 407–409.

Siekhaus, D.E., and Drubin, D.G. (2003). Spontaneous receptor-independent heterotrimeric G-protein signalling in an RGS mutant. Nat Cell Biol 5, 231–235.

Smith, J.A., and Rose, M.D. (2016). Kel1p Mediates Yeast Cell Fusion Through a Fus2p- and Cdc42p-Dependent Mechanism. Genetics 202, 1421–1435.

Tong, A.H., Evangelista, M., Parsons, A.B., Xu, H., Bader, G.D., Page, N., Robinson, M., Raghibizadeh, S., Hogue, C.W., Bussey, H., et al. (2001). Systematic genetic analysis with ordered arrays of yeast deletion mutants. Science 294, 2364–2368.

van Drogen, F., Mishra, R., Rudolf, F., Walczak, M.J., Lee, S.S., Reiter, W., Hegemann, B., Pelet, S., Dohnal, I., Binolfi, A., et al. (2019). Mechanical stress impairs pheromone signaling via Pkc1-mediated regulation of the MAPK scaffold Ste5. J Cell Biol 218, 3117–3133.

Volfson, D., Marciniak, J., Blake, W.J., Ostroff, N., Tsimring, L.S., and Hasty, J. (2006). Origins of extrinsic variability in eukaryotic gene expression. Nature 439, 861–864.

Wagih, O., Usaj, M., Baryshnikova, A., VanderSluis, B., Kuzmin, E., Costanzo, M., Myers, C.L., Andrews, B.J., Boone, C.M., and Parts, L. (2013). SGAtools: one-stop analysis and visualization of array-based genetic interaction screens. Nucleic Acids Res 41, W591–596.

Wang, Z., and Zhang, J. (2011). Impact of gene expression noise on organismal fitness and the efficacy of natural selection. Proc Natl Acad Sci U S A 108, E67–76.

Yu, L., Qi, M., Sheff, M.A., and Elion, E.A. (2008). Counteractive control of polarized morphogenesis during mating by mitogen-activated protein kinase Fus3 and G1 cyclin-dependent kinase. Mol Biol Cell 19, 1739–1752.

Zarzov, P., Mazzoni, C., and Mann, C. (1996). The SLT2(MPK1) MAP kinase is activated during periods of polarized cell growth in yeast. EMBO J 15, 83–91.

Zhang, N.N., Dudgeon, D.D., Paliwal, S., Levchenko, A., Grote, E., and Cunningham, K.W. (2006). Multiple signaling pathways regulate yeast cell death during the response to mating pheromones. Mol Biol Cell 17, 3409–3422.

Zhu, J., Deng, S., Lu, P., Bu, W., Li, T., Yu, L., and Xie, Z. (2016). The Ccl1-Kin28 kinase complex regulates autophagy under nitrogen starvation. J Cell Sci 129, 135–144.

